# Bottom-up reconstruction of functional Death Domain Signalosomes reveals a requirement for polymer stability and avidity

**DOI:** 10.1101/2024.05.08.593169

**Authors:** Mauriz A. Lichtenstein, Fakun Cao, Finn Lobnow, Paulina Dirvanskyte, Anna Kulesza, Elke Ziska, Marcus J. Taylor

**Author notes:** These authors contributed equally (Mauriz A. Lichtenstein, Fakun Cao, Finn Lobnow).

## Abstract

A key feature of innate immune signaling is the compartmentalization of signaling effectors into cellular structures referred to as signalosomes. Critical to the formation of these compartments are protein polymers composed of Death Domains (DD). However, the biophysical properties these polymeric scaffolds require for signal transduction are not clearly defined. Here, we engineered a single-component signalosome, referred to as Chimeric Higher-order Assemblies for Receptor Mediated Signaling (CHARMS). We found that CHARMS functionality depends on the stability provided by the DD polymer, which could also be achieved with bacterial DDs and synthetic filament-forming domains. This demonstrates the importance of kinetic stability and inducibility, irrespective of the origin of the motif. By varying the multiplicity of TRAF6 interaction motifs, we demonstrate that avidity is a tunable property that can control the amplitude of signaling outputs. This work lays out a reductionist framework to dissect the required properties of signaling through polymeric scaffolds by adjusting their assembly kinetics, stability and avidity.

## Introduction

As cells navigate the informational content of their environment, accurately interpreting extracellular signals and transducing them intracellularly is critical for survival. A recurrent regulatory theme in signal transduction is the compartmentalization of biochemical reactions into multi-protein complexes termed signalosomes (*1–3*). To understand how cells sense and respond to chemical inputs we need to understand how signalosomes assemble, recruit, and activate effector proteins. A striking example within signalosomes in innate immune signaling is the use of molecular scaffolds: upon a trigger, protein monomers in solution assemble into filaments (*3*, *4*). The polymeric filaments at the core of these signalosomes are composed of death domain (DD) superfamily proteins or Toll/IL-1R (TIR) domain-containing proteins (*5*, *6*), two ancient protein domains found in both bacteria and eukaryotes (*7*, *8*). The signaling output of these signalosomes is determined by the effector proteins recruited to this scaffold, as the oligomers form concentrated foci of effector binding sites. Many DD-containing proteins have binding motifs for the E3 Ligases of the TRAF family (*9*), and TRAF6 activation serves as a convergence point for multiple innate immune signaling scaffolds culminating in inflammatory NF-kB signaling. These unifying structural features suggest these signaling scaffolds at the core of signalosomes have a common signal transduction mechanism.

If innate immune oligomeric scaffolds have a shared signaling mechanism, a core set of biophysical properties likely governs their signaling output. However, despite shared mechanistic features, innate immune signaling scaffolds are diverse: while signalosomes commonly feature open-ended filamentous assemblies, such as in the Rig-1-like Receptor complex (*10*) or the Card11-Bcl10-Malt1 complex (*11*), others feature stoichiometrically defined oligomers, such as in the Myddosome. In addition, they can consist of variable combinations of DDs or TIR domain-containing proteins and function from distinct cellular locations (*4*, *12*, *13*). This diversity has made it difficult to define the necessary biophysical properties and the minimal number of components required to assemble a functional scaffold. Over the last decades, detailed mechanisms of signalosome function were determined by mutational or interaction studies. These top-down approaches can characterize the biochemical and cell biological diversity that operate at individual signalosomes. However, to determine how these signalosomes converge on TRAF6 and NF-kB signaling, we need to define unifying design principles spanning all signaling scaffolds. This requires a bottom-up approach - building a simplified signaling system that can be used to reconstruct the behavior of signaling scaffolds *de novo*.

### A single chimeric protein can functionally replace a three-component innate immune signalosome

As protein domains encode functional properties, by rearranging these domains and creating a minimal functioning signaling unit we can identify which core properties are needed for signalosome function. We aimed to do this by reducing a multi protein signalosome to its core components by creating chimeric fusion proteins. For our model system, we chose the Myddosome - a three-component signalosome composed of MyD88, IRAK4 and IRAK1 proteins that form a filamentous scaffold upon the triggering of Toll-like and interleukin-1 (IL-1) receptors (*14*, *15*). TRAF6 is recruited to the Myddosome via TRAF6 binding motifs (T6BM) found in IRAK1 (Fig. 1A), sharing the amino acid sequence PxExxZ (*9*) (where Z is an aromatic or acidic residue). Ultimately, this leads to pro-inflammatory NF-kB signaling. To create an orthogonal reaction vessel to test the chimeric proteins, we created a triple-knockout (3xKO) cell line for MyD88, IRAK4 and IRAK1 in EL4 mouse lymphoma cells, that also expressed TRAF6 tagged with mScarlet (fig. S1A). The 3xKO cells cannot signal in response to IL-1 stimulation (Fig. S1B), hence any restoration of IL-1 signaling activation upon expression of chimeric proteins would show reconstitution of a functioning signaling pathway. To determine the unifying design principles spanning all innate immune signaling scaffolds, we designed a system restricted to their common features: the ability to read out information from receptors, the presence of DDs and TRAF6 binding motifs (T6BMs).

**Figure 1:**
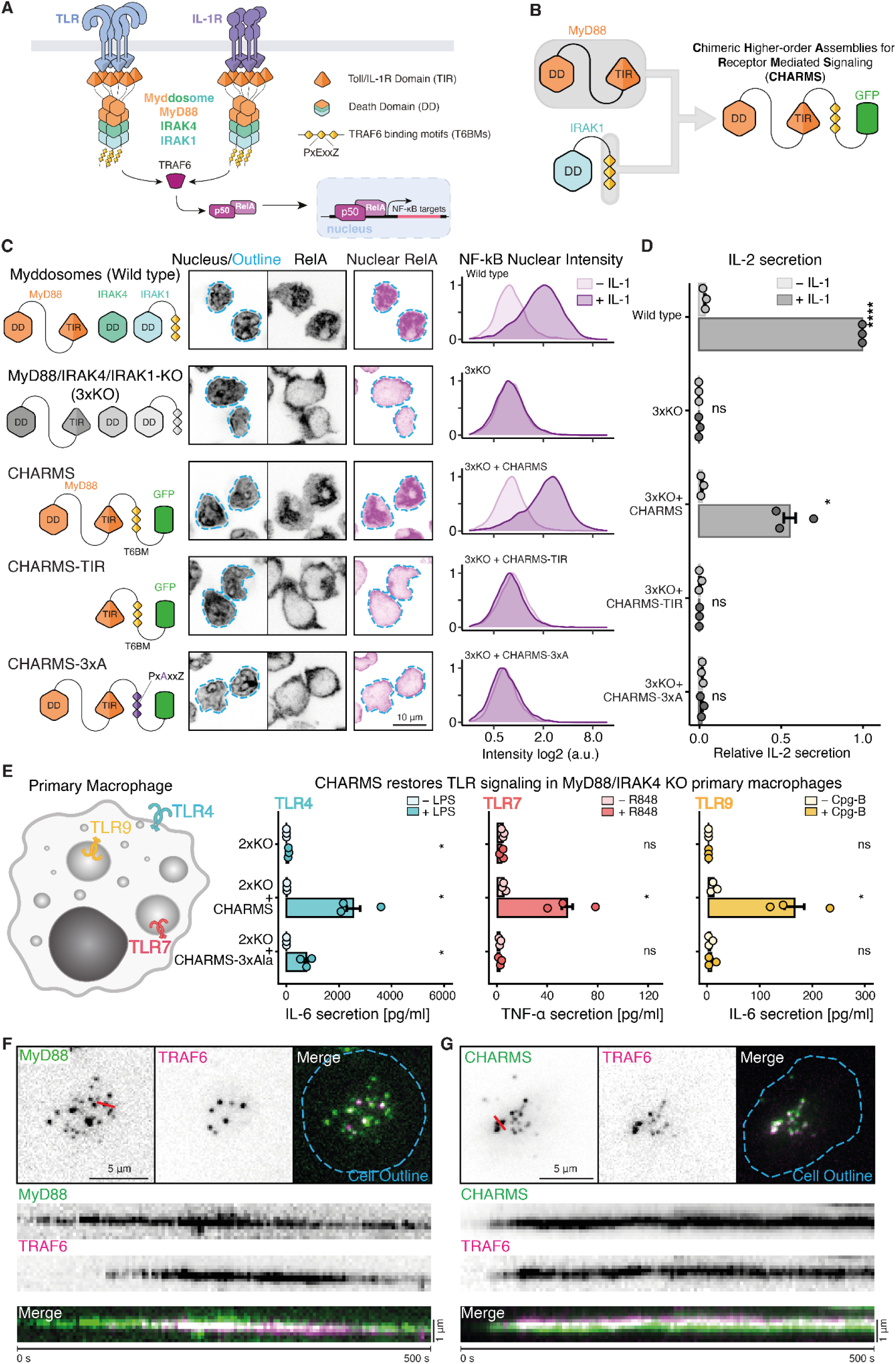
Single component CHARMS reconstructs the behavior and signaling output of a three component innate immune signalosome. **(A)** The Myddosome is a three component signalosome that assembles and transduces signals in response to TLR and IL-1Rs stimulation. Critical to Myddosome functionality are the TIR domain, Death domain, and TRAF6 binding motifs. The formation of this signalosome leads to the recruitment and activation of TRAF6 resulting in NF-kB (p50/RelA) translocation to the nucleus. **(B)** To determine the design principles of innate immune signaling signalsomes, we designed a chimeric fusion protein consisting of full length MyD88 fused to the C-terminus of IRAK1 that contains 3xT6BMs (PxExxZ). We called this system Chimeric Higher-order Assemblies for Receptor Mediated Signaling (CHARMS). **(C)** Restoration of NF-kB activation in response to IL-1 in 3xKO expressing CHARMS. Left, schematic showing the different CHARMS variants tested in 3xKO cells. Middle, confocal micrographs showing the localization of RelA after IL-1 stimulation in WT, 3xKO cells, and 3xKO cells expressing CHARMSs variants. Dashed cyan outline superimposed on the Hoechst images shows segmented nuclear outline. This was used to segment and measure the RelA nuclear staining (third column). Scale bar, 10 µm. Right, quantification of the RelA nucleus intensity (normalized to cytoplasmic RelA staining) in unstimulated and IL-1 stimulated cells. Distributions of RelA nuclear intensity composed of measurement taken from N= 1138 - 7789 cells. **(D)** Restoration of cytokine release in response to IL-1 stimulation in 3xKO cells expressing CHARMS. ELISA measures IL-2 release in WT and 3xKO cells expressing CHARMS variants. IL2 release was normalized to that of stimulated WT cells and baseline-corrected using unstimulated 3xKO. The p-values are **** =0.0001 and * =0.01. Bars represent mean ± s.e.m (N = 3 biological replicates measured per cell line). Statistical significance is determined using an unpaired t-test. **(E)** Reconstitution of CHARMS in primary macrophages derived from MyD88-/-,IRAK4-/- (2xKO) mice restores TLR signaling from plasma membrane and intracellular membranes. Schematic depicts primary macrophages with the subcellular localization of the TLRs tested. Plots show IL-6 or TNF-α release after stimulation with TLR4, 7 or 9 agonists. TLR4 was stimulated with 100 ng/ml LPS for 4h and TLR9 was stimulated with 1 µM Cpg-B (ODN1668) for 6h to measure IL-6 secretion. TLR7 was stimulated with 50 ng/ml Resiquimod (R848) for 8h to measure TNF-α secretion. Statistical significance between unstimulated and stimulated conditions determined using unpaired t-test. The p-values are * < 0.05. Bars represent mean ± s.e.m (N = 3-4 independent biological replicates measured per cell line, n = 3 experimental replicates performed within the same experiment). **(F-G)** IL-1 triggered assembly of CHARMS into plasma membrane associated puncta that recruit TRAF6. TIRF images showing the recruitment of mScarlet-TRAF6 to WT Myddosomes (right) and CHARMS assemblies in cells undergoing IL-1 stimulation, Scale bar, 5 µm. Kymograph analysis shows an example of Myddosome formation (left) and CHARMS assembly (right) that recruit TRAF6. Kymograph taken from the red line overlaid MyD88/CHARMS images. Scale bar, 2 µm.

We created three chimeric fusion proteins containing all or some of these features. First, we fused full length MyD88 to the C-terminus of IRAK1 that contains 3xT6BMs (PxExxZ), creating a single chimeric MyD88 protein (Fig. 1B). We called this system **C**himeric **H**igher-order **A**ssemblies for **R**eceptor **M**ediated **S**ignaling (CHARMS). To test the minimal sufficient scaffold that can activate TRAF6 and NF-kB signaling we created two further chimeric proteins (CHARMS-TIR, CHARMS-3xA). A construct consisting of the TIR domain of MyD88 fused to the T6BMs of IRAK1 (CHARMS-TIR, Fig. 1C, *middle*) and a version of chimeric MyD88 with the central glutamic acids of T6BM mutated to alanine (CHARMS-3xA, Fig. 1C, *bottom*) that abolishes the interaction with TRAF6 (*9*). We expressed CHARMS in 3xKO cells and found it could qualitatively rescue signaling, restoring the relocation of cytoplasmic RelA to the nucleus upon IL-1 stimulation (Fig. 1C, fig. S1C). Conversely, CHARMS-TIR or CHARMS-3xA cannot rescue RelA translocation (Fig. 1C, Fig. S1C). Only CHARMS restored downstream signaling output as measured by cytokine release (Fig. 1D), suggesting that successful signaling depends on the DD and the recruitment of TRAF6 via T6BM. This shows that a single chimeric fusion protein, containing just the TIR and DD of MyD88 and T6BM of IRAK1 contains the necessary properties required for IL-1 signaling. This system allows us to test and determine the essential properties of a signalosome required to activate TRAF6 and NF-kB.

In addition to IL-1Rs, TLRs can signal through Myddosomes, owing to the homotypic interactions of the TIR domains of MyD88 and TIR domains of the TLRs (*16*, *17*). To test whether the CHARMS can signal downstream of the TLRs, we generated bone marrow derived macrophages from a MyD88/IRAK4 double KO (2xKO) mouse (fig. S1D), reconstituted these cells with CHARMS and tested whether it restored signaling responses to three TLR ligands. The chimeric signalosome restores signaling in the 2xKO cells in response to the triggering of plasma membrane associated TLR4, and two TLRs that signal from intracellular membranes TLR7 and TLR9 (*18*) (Fig. 1E). With this experiment we can establish that our chimeric signalosome retains a key property of Myddosomes - the ability to read out different receptors by interacting with TIR domains of both IL-1R and TLRs. This also shows that CHARMS can transduce signals from diverse cellular locations with distinct biochemical environments.

In cells, signalosomes that assemble around oligomeric protein scaffolds form large cytoplasmic puncta that can be distinguished using fluorescence microscopy (*19*, *20*). To verify whether our chimeric proteins signal via the formation of a signalosome-like protein scaffold near the cell membrane, we used total internal reflection fluorescence microscopy (TIRFM) and supported lipid bilayers (SLBs) functionalized with IL-1 (*21*). Upon IL-1 stimulation in a WT background, MyD88 forms puncta, which increase in intensity and stably recruit TRAF6 (Movie 1, Fig. 1F, fig. S1E). We only observe comparable molecular dynamics for GFP-tagged CHARMS with functional TRAF6 binding motifs (Movie 2, Fig. 1G, fig. S1E, F). Thus, CHARMS, a single-component chimeric MyD88, can functionally replace the three-component Myddosome in living cells. CHARMS can assemble in response to receptor stimulation into large signalosomes that have qualitatively similar properties and signaling output to Myddosomes. We conclude that CHARMS has the required biophysical properties to activate TRAF6, NF-kB and downstream transcriptional responses.

### Reconstruction of functional CHARMS signalosomes from diverse filament forming domains

For its signaling function, we hypothesize that CHARMS requires at least two properties – it must be able to oligomerize into a scaffold and recruit an effector. Importantly an essential criterion is that both these properties must be controlled by receptor activation (*22*). To investigate whether these properties are distinctly encoded by the different domains, we analyzed the role of the TIR and DD in assembling a functional signalosome more closely. The MyD88 TIR domain and DDs can both form filamentous polymers (*6*, *15*) (Fig. 2A), suggesting a degree of functional overlap. However, given that the DD is required for signal transduction (Fig. 1C,D), it remains unclear what its role would be in a functional signalosome that cannot be assumed by the TIR domain. To analyze this further we examined the ability of the CHARMS-TIR construct to recruit TRAF6 in live cells using TIRFM. CHARMS-TIR assembles into cell surface puncta in response to IL-1 stimulation, but recruits TRAF6 only transiently (Fig. 2B,C). These results suggest that the DD provides the sufficient properties to the signalosome scaffold that allows it to stably recruit TRAF6.

**Figure 2:**
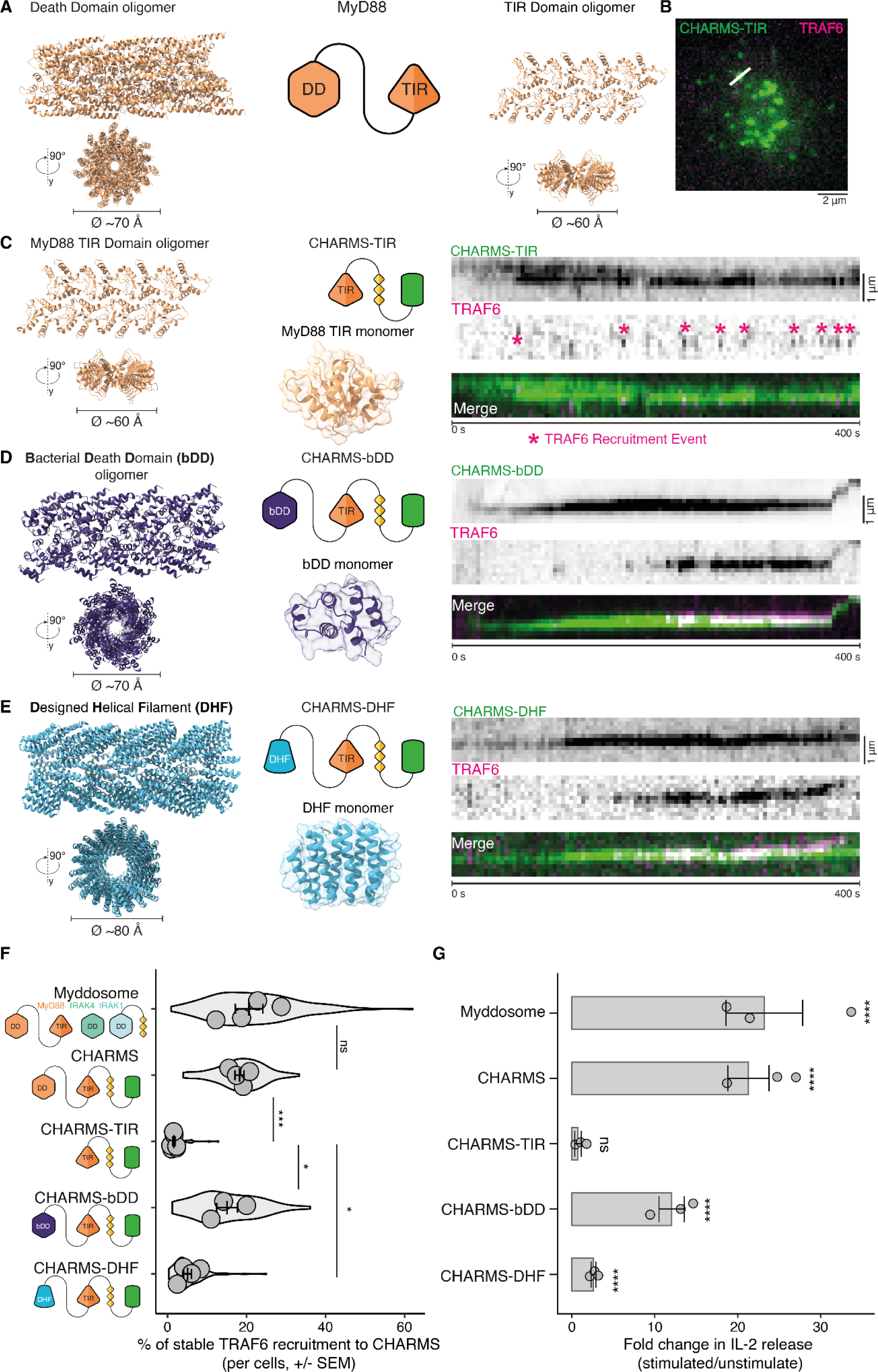
Functional replacement of the Death Domain by distantly related bacterial homologs or synthetic filament forming domains. **(A)** MyD88 is composed of two domains that can form filamentous polymers: the death domain and TIR domain. Shown are the solved helical filament MyD88-DD structure (PDB: 6I3N, helix extended using Rosetta) and the AF2-predicted filament of MyD88 TIR domains. **(B-E)** TIRF microscopy was used to analyze TRAF6 recruitment to CHARMS variants reconstituted in live cells. Shown is a TIRF image of the merged CHARMS-TIR and mScarlet-TRAF6 channels, the red line overlay corresponds to where the kymograph in (C) was taken. **(C)** CHARMS-TIR, which is solely composed of the MyD88-TIR domain fused to T6BMs, only transiently recruits TRAF6. Magenta asterisks on TRAF6 kymographs highlight TRAF6 recruitment events. Kymographs derived from red line overlaid TIRF images in (B). Scale bars, 5 µm. **(D)** CHARMS-bDD, a construct containing a bacterial DD (bDD) homolog from *Nostoc sp.* 106C (NCBI WP_086756477.1), restores stable TRAF6 recruitment. Left, AF2 prediction of the individual domain and two views of the predicted filamentous polymer. Right, kymograph analysis shows stable mScarlet-TRAF6 recruitment to CHARMS-bDD signalosomes in 3xKO cells. **(E)** CHARMS-DHF, a construct containing a synthetic filament forming domain ((*23*) PDB: 6E9X), restores stable TRAF6 recruitment. Left, structure view of the individual DHF domain and two views of the filamentous polymer. Right, kymograph analysis shows stable mScarlet-TRAF6 recruitment to CHARMS-DHF signalosomes in 3xKO cells. **(F)** Quantification of stable TRAF6 recruitment events (≥40s) to CHARMS signalosomes. Shown is the percentage of stable recruitment out of total TRAF6 recruitment events per cell. Violin plots show the distribution stable TRAF6 recruitment events per individual cell across replicates. Mean percentage of stable TRAF6 ± s.e.m.: 20.6±3.5%, 18.2±1.1%, 1.6±0.2%, 15.1±2.6%, 5.0±1.0% for WT, CHARMS, CHARMS-TIR, CHARMS-bDD, and CHARMS-DHF respectively. Data points superimposed on the violin plots show the mean of individual experimental replicates, n = 4, 4, 6 3, and 5 replicates for cells expressing Myddosomes, CHARMS, CHARMS-TIR, CHARMS-bDD and CHARMS-DHF. The P-value are * < 0.05 and *** < 0.001. Statistical significance is determined using unpaired t-test. **(G)** Reconstitution of CHARMS-bDD and CHARMS-DHF restores IL-2 release in 3xKO cells in response to IL-1 stimulation. ELISA measurments of IL-2 release in EL4-MyD88-GFP/mScarlet-TRAF6 cells expressing all Myddosome components (referred to as WT in the figure) and 3xKO cells expressing CHARMS variants. The IL-2 release is shown as a fold change in the amount of IL-2 released in stimulated vs unstimulated cells. WT, CHARMS and CHARMS-TIR ELISA data the same as presented Fig. 1C, and shown here for comparison to CHARMS-bDD and CHARMS-DHF. The P-value are **** <0.0001. Bars represent mean ± s.e.m (mean calculated from n=3 independent experimental replicates). Statistical significance is determined using unpaired t-test.

If the DD encoded a discrete biochemical property required for signaling, we would expect that it would be similar in distantly related members of the superfamily. We chose a bacterial death domain (bDD) that, based on AlphaFold Protein Complex predictions, forms helical filaments, similar to the DD of MyD88 (*15*) (Fig. 2A,D, fig. S2A-B). CHARMS-bDD forms puncta that can recruit TRAF6 for extended periods of time (Fig. 2D). These results show that a bacterial homologue of the DD contains the requisite properties for scaffold formation that allows it to be functionally interchangeable with a mammalian DD, suggesting a wider conservation across this ancient superfamily.

If the propensity to form helical polymers is conserved within members of the DD superfamily that assemble signalsomes (*5*), then we predict that this structural feature underpins the domain’s functionality. As such, unrelated protein domains that can form qualitatively similar helical structures in response to a stimulus should be able to functionally replace the DD in signalosomes. To test this, we selected a computationally designed de novo protein (*23*) that assembles into helical filaments (Fig. 2E). We find that replacing the DD with the *in silico* designed helical filament (DHF) forming domain restores stable TRAF6 recruitment in response to IL-1 stimulation (Fig. 2E). Direct comparison of all constructs shows that introducing a filament forming domain, either DD, bDD or DHF significantly increases the percentage of long-lived (≥40 s) recruitment events compared to CHARMS-TIR (Fig. 2F, fig. S2C-D). Restored long-lived TRAF6 recruitment corresponds to restored cytokine release in response to IL-1 (Fig. 2G, fig. S2E). Together, these results show that CHARMS composed of diverse filament-forming oligomers are sufficient for stable TRAF6 recruitment and signalosome mediated signal transduction. Therefore filament-forming oligomers must contain the requisite biophysical properties required for designing a functional signalosome.

### Stability is an emergent property of polymeric scaffolds and is a requirement for signal transduction

Although the TIR domain can form oligomeric assemblies in response to IL-1, these are not sufficient to recruit TRAF6 and transduce signaling (Fig. 1C,D and Fig. 2C,F). As such, the DD, the bDD and DHF filament forming domains must share another property that emerges from their capacity to oligomerize and allow prolonged TRAF6 recruitment with downstream signaling. One shared property of the DHF and many structurally characterized DDs is the high polymer stability. This is possibly attributed to the similar helical oligomer structure where internal subunits, by virtue of having multiple buried interfaces, have lower off-rates and disassembly can only proceed from the ends (*5*, *15*, *23*). If this property were essential for signal transduction, we would expect it to be absent from the non-functional CHARMS-TIR that lacks a helical filament forming domain. We therefore used fluorescence recovery after photobleaching (FRAP) to compare protein turnover within these assemblies and found that the CHARMS, CHARMS-bDD and CHARMS-DHF formed stable assemblies with no monomer turnover; in comparison, the CHARMS-TIR had monomer turnover (Fig. 3A-E). We conclude that, alongside oligomerization and effector recruitment, an additional property required for signalosome functionality is polymer stability.

**Figure 3.**
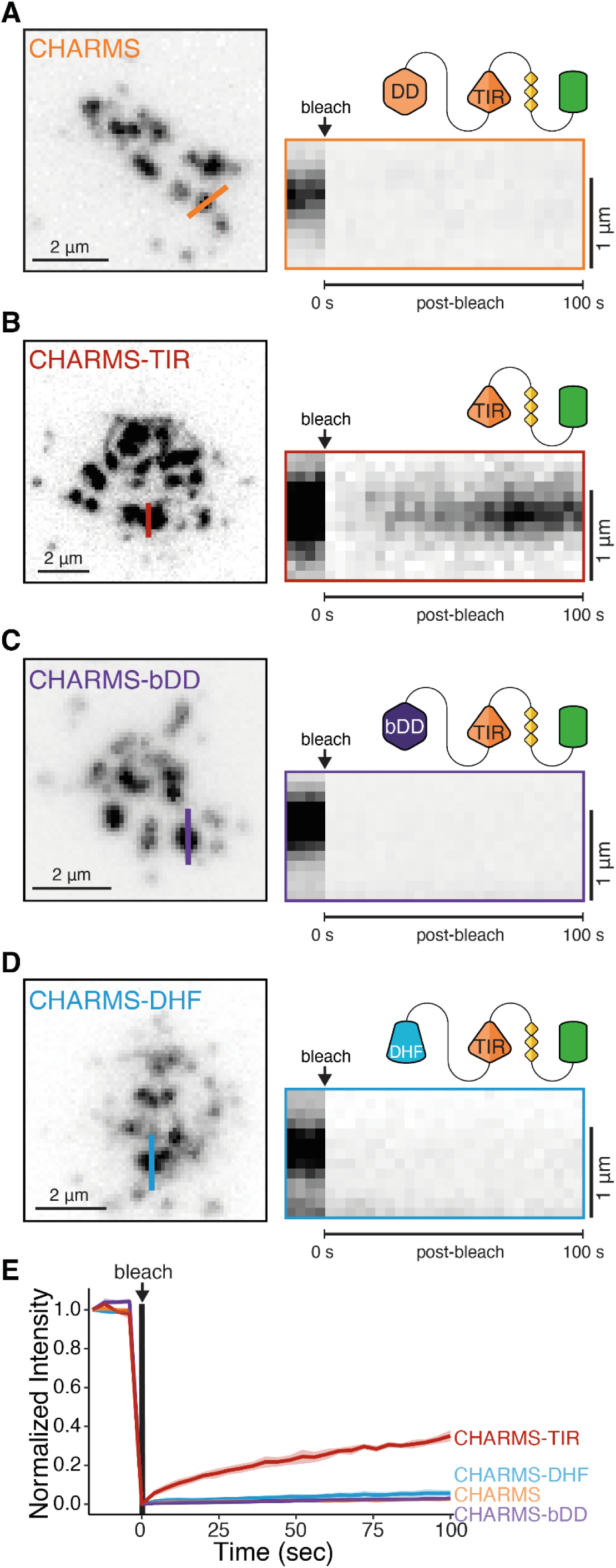
DDs or synthetic filaments provide high stability which is associated with functional CHARMS signalosomes. **(A-D)** FRAP analysis of CHARM variants. Left, TIRF images of the CHARMS signalosomes before photobleaching. The overlaid orange line on the TIRF images denotes the CHARMS assembly subjected to photobleaching. Right, kymograph analysis of the photobleached region. Kymographs derived from coloured lines overlaid TIRF images (left panel). Scale bars, 2 μm. **(E)** The quantification of the FRAP data shows recovery of CHARMS-TIR but not of other constructs. Lines represent the mean of experimental replicates and the shaded area corresponds to ± s.e.m. Measurements are taken from 3 independent replicates with N ≥9 regions of interest photobleached per replicate.

### Signaling output can be titrated by changing the number of effector recruiting motifs

Given that signaling output is a tunable response across many signaling systems (*24*), we next investigated which properties could achieve this. One property that could control signaling output is the density of T6BMs on the CHARMS scaffolds. The number of T6BMs varies on signaling effectors, for example IRAK1, IRAK2 or IRAK3 each possess three, two or one T6BM respectively (*9*). Because TRAF6 multimerization is required for further signal transduction (*25*), the number of T6BMs potentially controls the cooperative oligomerization and size of TRAF6 multimers. Thus signalosomes might naturally tune the connection to TRAF6 and thereby change how inputs are amplified. So far each designed CHARMS monomer has three effector binding sites (Fig. 1C and Fig. 2), and this raises the question whether the signaling output could be modulated by lowering or increasing the T6BMs multiplicity on the monomer (Fig. 4A). At the molecular level, the probability of TRAF6 recruitment increases with CHARMS signalosome size (fig. S3). Therefore, by changing the ability of the signalosome to recruit TRAF6, we could then change how the signalosome converts and amplifies the input from the receptor into a downstream response (Fig. 4A). To test this, we created a synthetic TRAF6 effector recruiting domain containing 1x, 3x or 5x T6BMs (Fig. 4B, fig. S4). All of these constructs form puncta and recruit TRAF6 (Fig. 4B); however, the TRAF6 recruitment to the CHARMS was modulated by the multiplicity of T6BMs. Increased multiplicity of binding motifs resulted in the formation of larger TRAF6 multimers associated with the CHARMS scaffold (Fig. 4C, fig. S5A,B); it also increased the stability of TRAF6 recruitment (Fig. 4D, fig. S5C,D). In addition, the probability of TRAF6 recruitment to CHARMS signalosomes increased with T6BM multiplicity (Fig. 4E, fig. S6A,B). Finally, this ability to engineer TRAF6 recruitment by increasing the multiplicity of T6BMS corresponded to elevated cytokine secretion (Fig. 4F, fig. S6C). This shows that the signaling output can be amplified by increasing the number of effector binding motifs on the scaffolds. We conclude that signaling output is an emergent property of the local concentration of effector binding sites within the signalosome.

**Figure 4.**
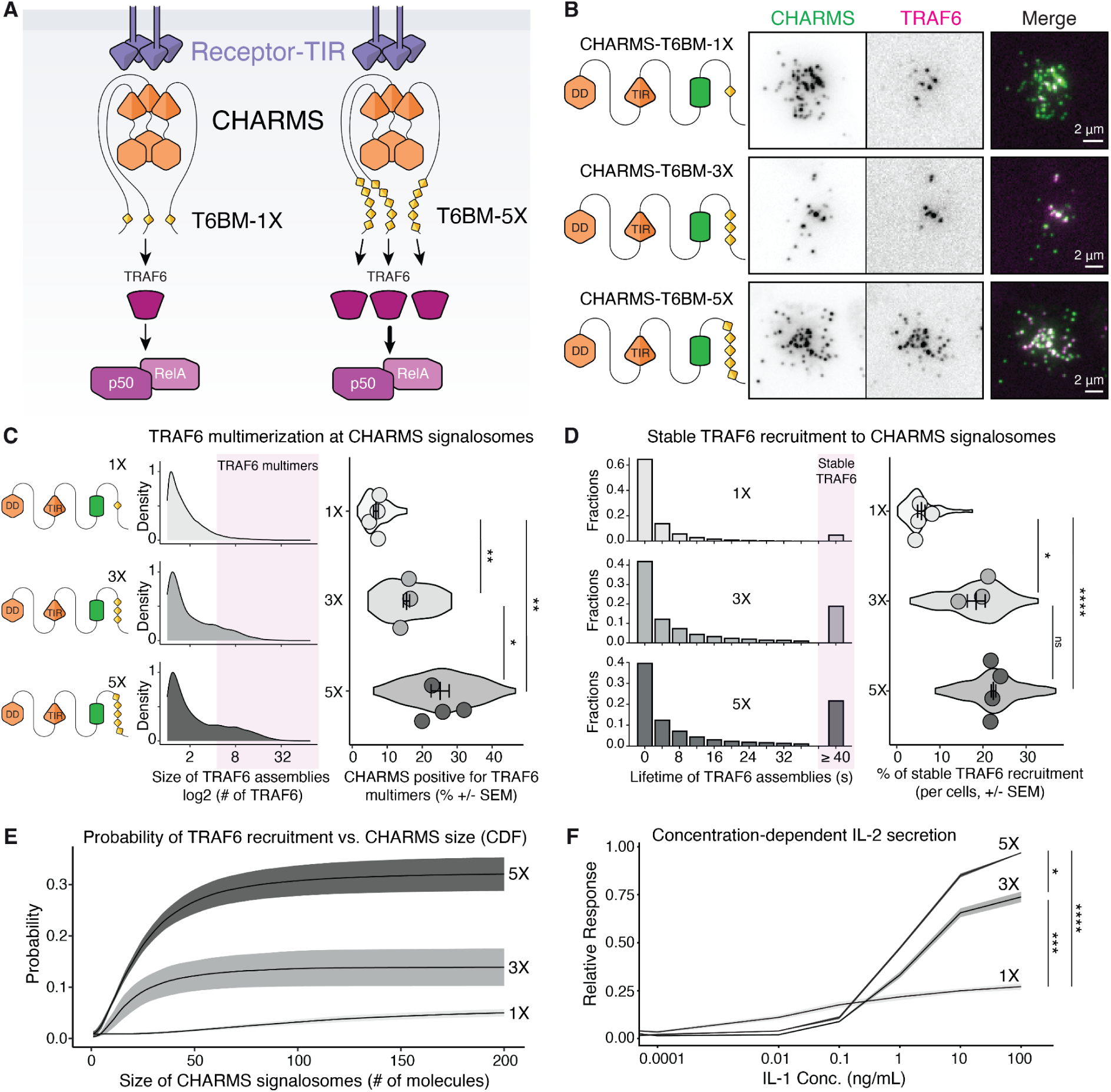
The avidity of binding sites to TRAF6 is tunable and can modulate signalosome properties and signaling output. **(A)** CHARMS connect to TRAF6 via TRAF6 binding motifs (T6BMs), this connection is required for TRAF6 activation and NF-kB signaling. We propose that varying the number and density of T6BMs on CHARMS scaffolds could change how the signalosome converts and amplifies the input from the receptor into a downstream response. **(B)** We designed CHARMS with 1, 3 and 5 T6BMs (PxExxZ) - CHARMS-1x (top), CHARMS-3x (middle) and CHARMS-5x (bottom), and reconstituted them into 3xKO cells. TIRF imaging confirmed that all three CHARMS variants could recruit TRAF6. Scale bar, 2 µm. **(C)** Greater multiplicity of T6BMs increases the size of TRAF6 assemblies associated with CHARMS. Left, schematic of CHARMS with 1, 3, and 5 T6BMs. Middle, Density plot of number of TRAF6 assemblies on CHARMS. Light magenta shaded area represents TRAF6 assemblies designated TRAF6 multimers (≥ 4.5 TRAF6 molecules). Right, quantification of the percentage of CHARMS signalosomes associated with TRAF6 multimers. Violin plots show the distribution of individual cell measurements. Colored dots superimposed on violin plots correspond to the average value per cell across independent experimental replicates. P values, * p < 0.05; ** p < 0.01. Quantifications are from n = 3-4 replicates with 6-20 cells measured per replicate. Bars represent mean ± S.E.M. Statistical significance is determined using unpaired t-test. **(D)** Multiplicity of T6BMs enhances the stability of TRAF6 at CHARMS signalosomes. Left, histograms show the lifetime distribution of TRAF6 recruitment to CHARMS signalosomes. Light magenta shaded area is the sum of all events with a lifetime greater than 40 sec, which represents stable TRAF6 association with CHARMS. Right, quantification of the percentage of long-lived (≥ 40s) recruitment out of all TRAF6 recruitment events. Violin plots show the distribution of individual cell measurements. Colored dots superimposed on violin plots correspond to the average value per cell across independent experimental replicates. P values, * p < 0.05; **** p < 0.0001. Quantifications are from n = 3-4 replicates with 12-27 cells measured per replicate. Bars represent mean ± s.e.m. Statistical significance is determined using unpaired t-test. **(E)** The multiplicity of T6BMs enhances the probability of TRAF6 recruitment. Cumulative distribution function (CDF) showing the probability of TRAF6 recruitment versus the size of CHARMS signalosomes. **(F)** The multiplicity of the TRAF6 binding motifs on CHARMS changes cytokine release. ELISA measuring of IL-2 release to an IL-1 stimulation dose-response. Gray-scale lines show the mean of three independent experimental replicates (N = 3). The shaded areas correspond to ±S.E.M. Statistical significance determined using one way ANOVA with Tukey correction. P values, * p < 0.05; *** p < 0.001 and **** p < 0.0001.

## Discussion

A major goal of biology is to be able to understand how functional biological systems emerge from their chemical constituents. Here we achieve this for innate immune signalosomes by using a bottom-up engineering approach. We engineered a chimeric fusion of MyD88 with TRAF6 binding motifs, referred to as CHARMS, and discovered this single protein fusion could functionally replace the three component Myddosome (Fig. 1). We found the death domain of CHARMS is critical to signal transduction, but can be functionally replaced by distant bacterial DD homologs or even a synthetic filament forming protein domain (Fig. 2). We define two key parameters for functionality. First, CHARMS require a high kinetic stability to recruit and nucleate TRAF6 assemblies capable of transducing downstream signals (Fig. 3). Second, the local density of TRAF6 binding motifs within CHARMS controlled the properties of TRAF6 recruitment, and presented a way to control downstream signaling output (Fig. 4). Like many innate immune signalosomes, the CHARMS is composed of a death domain oligomer and transduces signal via TRAF6 to the NF-kB pathway (*4*). Therefore the biophysical and biochemical parameters we identify as critical to CHARMS function may operate across a range of innate immune signaling cascades.

We show that a bacterial death domain can transduce information in a mammalian innate immune signaling pathway. This supports the idea that Death Domains have the requisite properties for accurate and inducible signaling, a key requirement for immune signaling which must detect and respond to threats to cell viability. This possibly explains the frequency of death domains in innate immune signalosomes (*5*, *26*). This is also consistent with the discovery of bacterial death domains in putative phage defense systems (*8*). However, we also show that synthetic proteins that assemble into qualitatively similar helical filaments can restore functionality as well. This suggests that the properties of polymer stability and inducible assembly that allow death domains to function as information transducers are possibly not unique. This is possibly the case for RIP homotypic interaction motif containing protein, which can assemble into amyloid (*27*), thus possibly serving as a polymeric scaffold that localizes downstream effectors . It is likely that unrelated protein folds might have acquired equivalent properties through convergent evolution. It also raises the possibility that these properties could be created synthetically through directed evolution approaches, opening up the ability to construct synthetic or orthogonal signalosomes.

We find that the signaling output of CHARMS can be controlled by varying the multiplicity of T6BMs (Fig. 4). It has been proposed that death domain oligomers function as high avidity scaffolds that concentrate signaling effectors, thereby compartmentalizing them within a signalosome (*4*, *5*). Our results support this model. Using a bottom-up, synthetic biology approach, we now demonstrate that avidity itself is tunable and can change a fundamental emergent property of a signaling cascade: the ability to convert and amplify an input into a downstream output. Critically, we show that avidity changes to the degree to which a given input is amplified. This finding might explain why the number of T6BM varies across different signaling effectors (*9*).

## Acknowledgements

We thank the staff at the Advanced Medical Bioimaging Core Facility, Charité Universitätsmedizin Berlin, for help with FRAP data acquisition and spinning disk confocal microscopy. We thank Uwe Klemm, Manuela Primke, Daniela Groine, and Bärbel Raupach for assistance and advice on breeding transgenic mouse lines; Arturo Zychlinsky for providing the MyD88 KO mouse line; Olivia Majer for advice on working with primary mouse macrophages and TLR stimulation experiments; Radhika Patnala and Arne Fabritius for advice on figure design and scientific illustrations; Lillian Fritz-Laylin for comments on figure design and layout; Simone Reber, Daniel Simpson and Randal Halfmann for critical reading and comments on the manuscript. We thank Iain Patten for assistance in drafting and constructing the manuscript. We are indebted to Kathrin Lattig for genotyping of transgenic mice and assistance with ELISA assays.

## Author contributions

M.J.T. conceived and supervised the study, designed and cloned the initial CHARMS constructs. E.Z. characterized the 3xKO cell lines, performed experiments in bone marrow progenitor cells from 2xKO mice with CHARMS, performed ELISA assays of TLR signaling in primary macrophages, and performed initial characterization of CHARMS in 3xKO cells. M.A.L. characterized the gene-edited 3xKO cell line expressing mScarlet-TRAF6, reconstituted and engineered 3xKO cells expressing CHARMS, designed CHARMS variants, performed ELISA, FRAP and microscopy analysis of CHARMS-expressing cells. F.L. performed in silico structural analysis of bacterial death domain homologs, and conducted ELISA and microscopy analysis with CHARMS-bDD. F.C. performed RelA staining and confocal microscopy analysis of RelA nuclear localization, conducted FRAP analysis of CHARMS, and performed microscopy analysis of CHARMS variants. P.D. isolated 2xKO bone marrow progenitor cells, performed transient immortalization, and performed collection of FRAP and microscopy data. A.K. performed IL-1 dose responses to compare CHARMS with different multiplicities of T6BMs. M.A.L., F.C., F.L., P.D. performed data analysis and visualization. M.J.T., M.A.L., F.C., F.L., E.Z. and P.D. wrote the manuscript.

## Supplemental Material

## Material and Methods

### Reagents TABLE

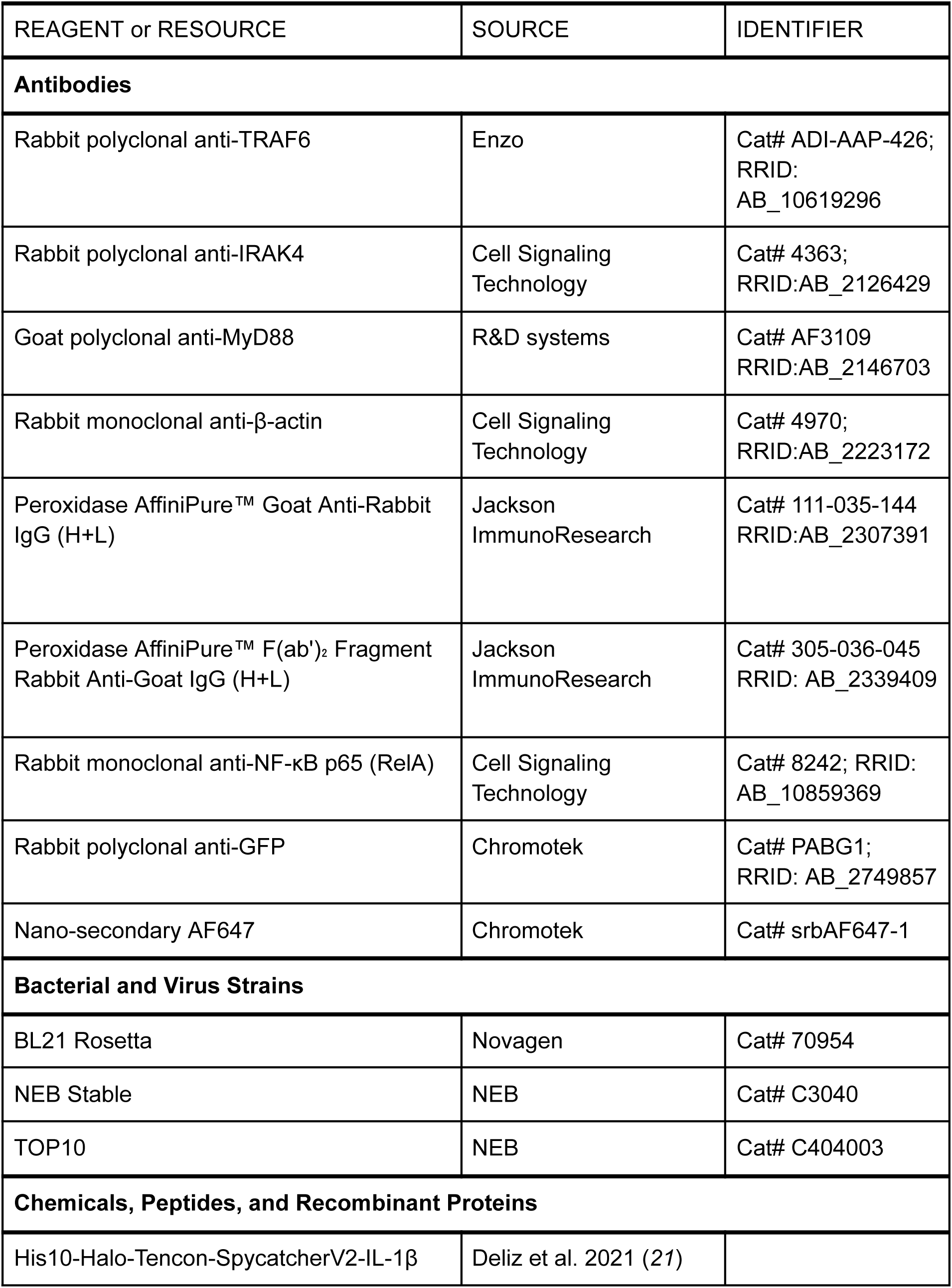
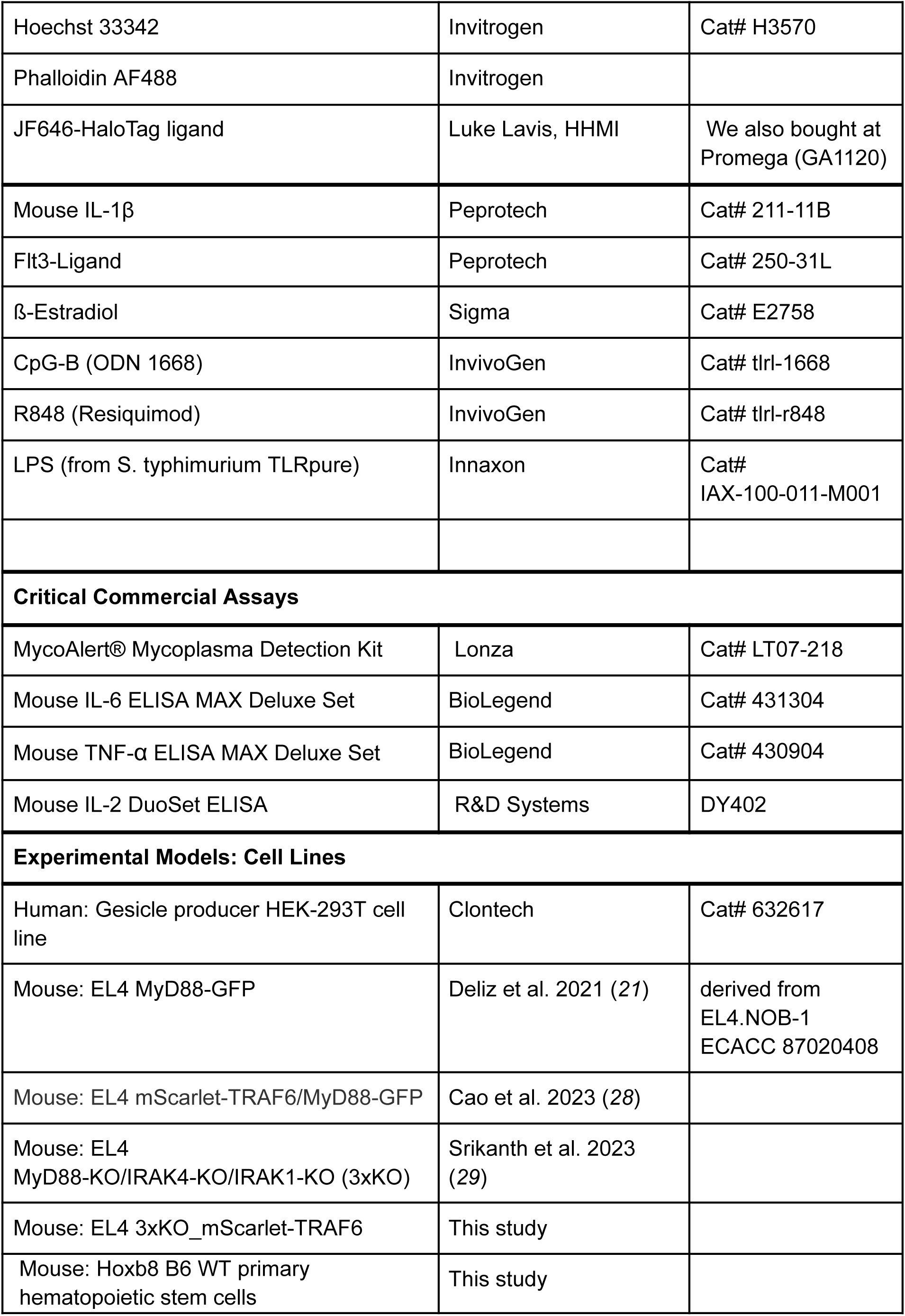
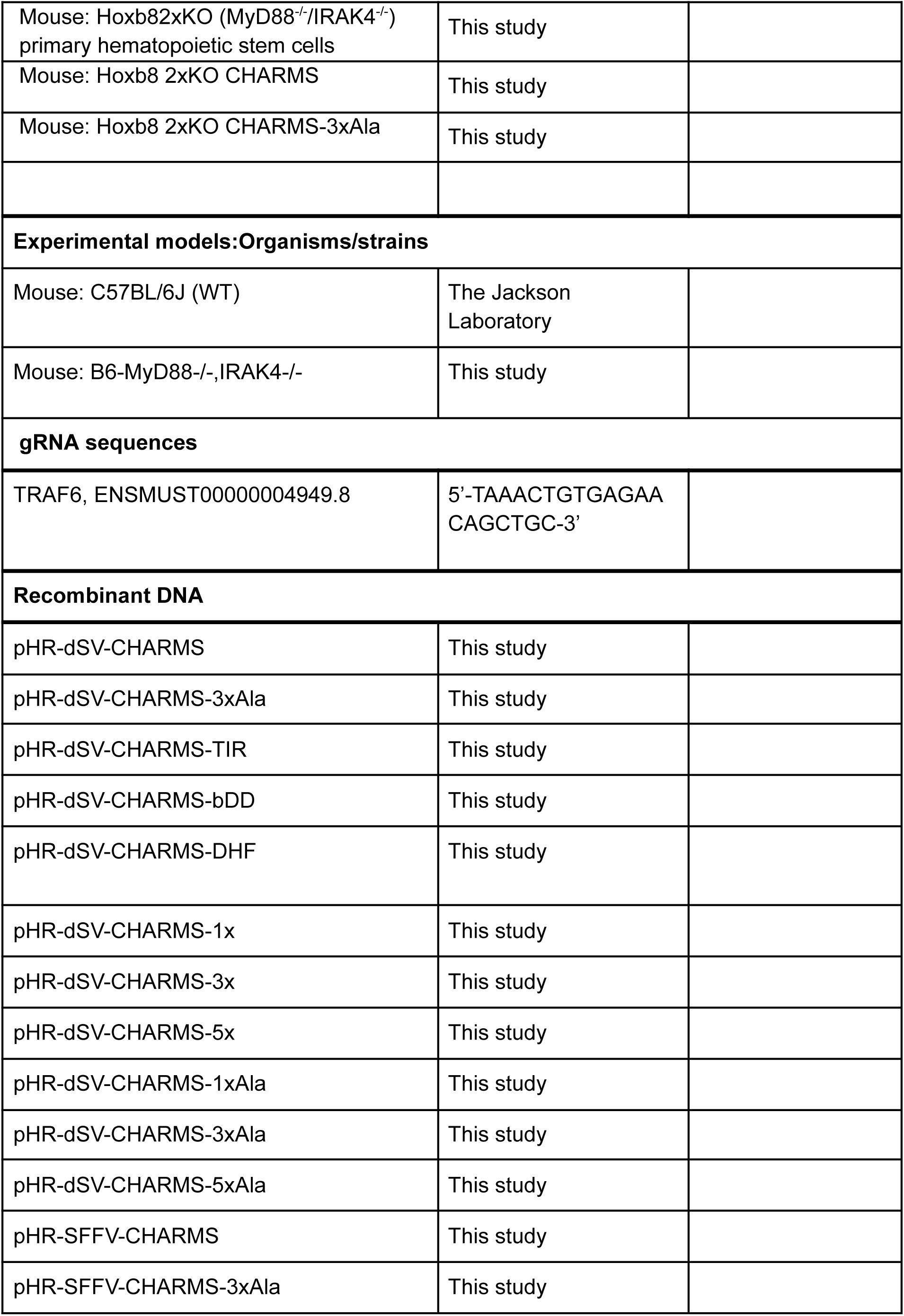
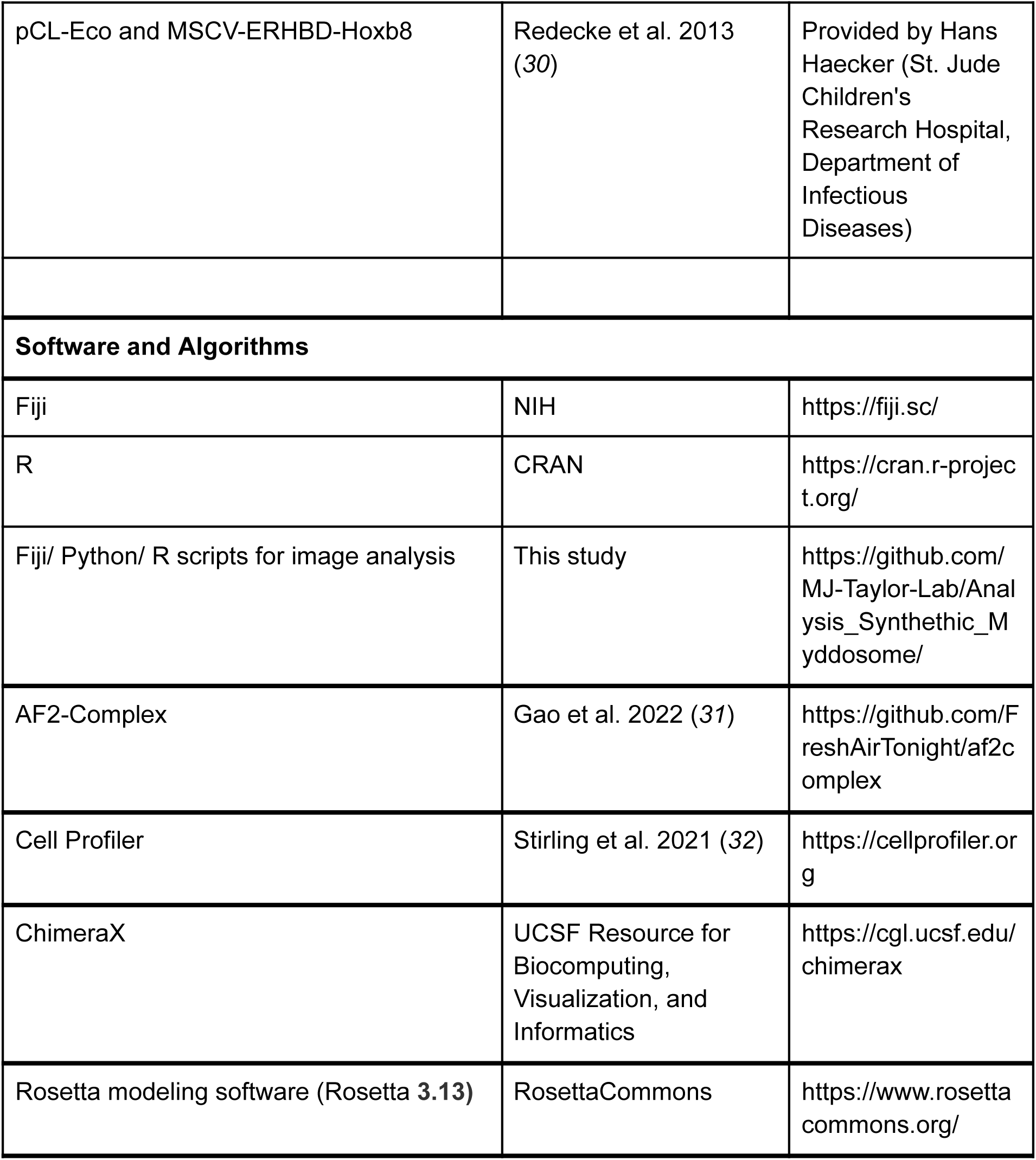

### Methods Cell Culture

All EL4 cell lines used in this paper are originally derived from EL4.NOB1 WT cells (ECACC, and referred to as EL4 in the paper) and were grown in RPMI (Thermo Fisher Scientific) with 10% FBS (Sigma) supplemented with 2 mM L-glutamine. EL4-3xKO cells were previously described (*29*). The EL4-MyD88-GFP/mScarlet-TRAF6 cell line was previously described (*28*). In the manuscript we used EL4-MyD88-GFP/mScarlet-TRAF6 cell line to compare Myddosomes to CHARMS, and is therefore referred to as WT in the figure legends and text (Fig. 1 and fig. S1). EL4 cultures were maintained at a cell density of 0.1-0.5x 10^6^ cells/ml in 5% CO2, 37°C. HEK-293T cells (Clontech) were grown in DMEM (Thermo Fisher Scientific) with 10% FBS supplemented and 2 mM L-glutamine. All cells were determined to be negative for mycoplasma using the MycoAlert detection kit (Lonza).

### Differentiation and culture of bone marrow derived primary mouse macrophage cells

Macrophages were differentiated from bone marrow derived hematopoietic/myeloid precursors that were transiently immortalized. Bone marrow cells originated from male or female B6 (WT) or B6-MyD88-/-,IRAK4-/- (2xKO MyD88/IRAK4) mice. (2xKO mice were bred by crossing previously described IRAK4 (*33*) and MyD88 (*34*) KO mouse strain, final genotype: B6.129P2-Irak4^tm1Yeh^Cnbc-Myd88^tm1Aki^/J). For transient immortalization of hematopoietic/myeloid precursor cells the bone marrow of 6-12-wk-old mice was isolated transduced with an estrogen-regulated allele of Hoxb8 by retroviral delivery (*30*). Transient immortalized hematopoietic/myeloid precursors cells were maintained in RP10 progenitor medium, a formulation of RPMI 1640 medium, 10 % fetal calf serum, 1 % L-Glutamine, 1 % Sodium-Pyruvate, 1 % HEPES, 1 % Penicillin/Streptomycin with Flt3-Ligand (70 ng/ml, Peprotech) and ß-Estradiol (1µM, Sigma). The precursor cells were differentiated to macrophages in RP10 differentiation medium (RPMI 1640 medium, 10 % fetal calf serum, 1 % L-Glutamine, 1 % Sodium-Pyruvate, 1 % HEPES, 1 % Penicillin/Streptomycin, 10 % conditioned M-CSF medium produced by a 3T3 mouse embryonic fibroblast cell line, and 0,0003 % β-mercaptoethanol) in non-TC treated cell culture dishes. On day 7 differentiated 2xKO and 2xKO expressing CHARMS or CHARMS-3xAla macrophages were re-seeded into new cell culture plates for stimulation experiments on the following day. WT derived macrophages were included in all experiments as a positive control.

### Generation of CRISPR/Cas9 engineered cell lines

#### Endogenous labeling of TRAF6 with mScarlet in EL4-3xKO cells using CRISPR/Cas9

We purchased EL4 3xKO cells (*29*) that had been electroporated with Cas9 RNPs loaded a sgRNA targeting the TRAF6 gene locus (gRNA sequence 5’-TAAACTGTGAGAACAGCTGC-3’, ENSMUST00000004949.8) and a ssDNA homology direct repair template from the Synthego Corporation (Redwood CA, USA). From this pooled population, we isolated single cell clones by limited dilution on 96 well plates. We screened and identified single cell clones where a mScarlet-i ORF was inserted after the start codon of TRAF6 creating an inframe fusion. Positive clones were identified by western blot analysis (Fig. S1A), performed with the following antibodies: rabbit polyclonal anti-TRAF6 (Enzo, # ADI-AAP-426 and Rabbit monoclonal anti-β-actin (CST, # 4970)

### Generation of lentiviral constructs

Full details of the Lentiviral plasmid construction are given below. All full length constructs were confirmed by Sanger sequencing. The amino-acid sequences of all constructs are provided in Table S1, and the identifiers of the sequences (if applicable) in Table S2.

### pHR-dSV-CHARMS, pHR-dSV-CHARMS-3xAla, pHR-SFFV-CHARMS, pHR-SFFV-CHARMS-3xAla

The sequences encoding human IRAK1^538-712^ and the same sequence with E to A mutations in the critical PxExxZ motif (IRAK1^538-712/3xA^, E541A, E584A and E704A) were codon optimized for expression in mouse and ordered as gBlocks (IDT). These fragments were fused to a PCR fragment encoding codon optimized mouse MyD88 (amplified by PCR using the forward 5’-tttttggaggcctaggctacgcgtgccaccatgtctgcgggagacccccgcgtgggatc-3’ and 5’-ggacccgccactacctccgggcagggacaaagccttggcaaggcgggt-3’ reverse primers), mEGFP (amplified by PCR using the forward 5’-gggggtagtggaggaagtgtgagcaagggcgaggagctgttc-3’ and reverse 5’-tgcaggtcgactctagagtcgcggccgcttta-3’ primers) and a Mlu1/Not1 digested pHR-dSV or pHR-SFFV lentiviral plasmid using Gibson assembly. MyD88 was fused to IRAK1^538-712^ or IRAK1^538-712/3xA^ via a 2xGGS linker. mEGFP was fused c-terminally via a GGSGGTGGSGGS linker.

### pHR-dSV-CHARMS-TIR

pHR-dSV-CHARMS was used as templates in a PCR reaction using the forward 5’-ttggaggcctaggctacgcgtgccaccatgccacaaacaaaggaactgggaggc-3’ and reverse 5’-tgcaggtcgactctagagtcgcggccgcttta-3’ primers. The fragment was cloned into a Mlu1/Not1 digested pHR-dSV lentiviral plasmid using Gibson assembly.

### pHR-dSV-CHARMS-bDD

The sequence encoding for amino acids 230-303 of the CHAT domain-containing protein of the Cyanobacteria Nostoc sp. 160 (identifier: WP_086756477.1) was codon optimized for expression in mouse and ordered as a gBlock (IDT). These amino acids have been previously annotated as encoding for a member of the Death-Domain superfamily, hence the abbreviation bDD (bacterial Death-Domain (*8*)). To amplify the sequence encoding for MyD88^126-296^-IRAK1^538-712^-mEGFP, pHR-dSV-MyD88-IRAK1^538-712^-mEGFP was used as a template in a PCR reaction using the forward 5’-agcctttacaggtggccagagtggaaagcagtg-3’ and reverse 5’-tgcaggtcgactctagagtcgcggccgcttta-3’ primers. The gBlock was cloned together with the PCR fragment into a Mlu1/Not1 digested pHR-dSV lentiviral plasmid using Gibson assembly. bDD was fused to MyD88^126-296^ via a 2xGGS linker.

## pHR-dSV-CHARMS-DHF

The protein sequence of designed helical filament protein (DHF91 from (*23*)) was codon optimized for expression in mice cells and ordered as a gBlock (IDT). To amplify the sequence encoding for MyD88^126-296^-IRAK1^538-712^-mEGFP, pHR-dSV-MyD88-IRAK1^538-712^-mEGFP was used as a template in a PCR reaction using the forward 5’-ggaggtagtggcgggtccgagaagcctttacaggtggc-3’ and reverse 5’-tgcaggtcgactctagagtcgcggccgcttta-3’ primers. The gBlock was cloned together with the PCR fragment into a Mlu1/Not1 digested pHR-dSV lentiviral plasmid using Gibson assembly. DHF91 was fused to MyD88^126-296^ via a 2xGGS linker.

### pHR-dSV-CHARMS-1x, pHR-CHARMS-3x, pHR-CHARMS-5x, pHR-dSV-CHARMS-1xAla, pHR-dSV-CHARMS-3xAla and pHR-dSV-CHARMS-5xAla

We chose the well characterized TRAF6-binding-motif (T6BM) of CD40^229-238^ (PKQEPQEIDF) (Ye et al. 2002). The T6BMs are separated by 15 amino acid long GGS linkers. The sequences encoding for 1x-T6BM, 3x-T6BM, 5x-T6BM, 1x-T6BM^E/A^, 3x-T6BM^E/A^ and 5x-T6BM^E/A^ were codon optimized for expression in mouse and ordered as a gBlock (IDT). To amplify the sequences encoding for MyD88-mEGFP, a previously described plasmid encoding for MyD88-GFP (*21*) was used as a template in a PCR reaction using the forward 5’-cttttttggaggcctaggctacgcgtgccaccatgtctgcgggag-3’ and reverse 5’-acccttgtacagctcgtccatgcc-3’ primers. The gBlock was cloned together with the PCR fragment into a Mlu1/Not1 digested pHR-dSV lentiviral plasmid using Gibson assembly.

### Lentiviral production and transduction of cell Lines

Lentivirus particles were produced in HEK-293T cells by co-transfection of the pHR transfer plasmids with second-generation packaging plasmids pMD2.G and psPAX2 (a gift from Didier Trono, Addgene plasmid # 12259 and # 12260). Virus particles were harvested from the supernatant after 48 hours, filtered with a 45 um spin filter unit, and applied to EL4 cells. 24 hours later fresh medium was added on top and another 24 hours later the medium was completely exchanged.

FACS was performed to select cell populations with the homogenous expression levels of CHARMS alleles. EL4-3xKO mScarlet-TRAF6 cells were used as a negative control and EL4-MyD88-GFP knock in cell lines (*21*) were used to create a sorting gate that selected lentivirus transduced cells with equivalent expression levels. For experiments that examined the multiplicity of TRAF6 binding motifs (Fig 4), we sorted homogenous cell populations with equivalent expression levels of CHARMS variants, thereby allowing us to quantitatively compare signalosomes composed CHARMS-1x, 3x and 5x and downstream signaling outcomes (fig. S4). All cell sorting was done using a BD FACS Aria II at the Deutsches Rheuma-Forschungszentrum Berlin Flow Cytometry Core Facility.

To transduce primary hematopoietic stem cells that had been transiently immortalized, we applied the same steps as described above with the following changes. 12h after co-transfection of transfer and packaging plasmids to the HEK-293T cells the medium was changed to RP10 medium w/o ß-Estradiol. Viral supernatant was collected after 48h, cleared by centrifugation and applied to primary hematopoietic progenitor cells by spin-infection in RP10 medium supplied with 4µg/ml Polybrene, 1µM ß-Estradiol and 70 ng/ml Flt3-Ligand. Infected cells were incubated overnight at 32°C. 48 hours later the medium was completely exchanged to the progenitor medium and cells were checked for GFP expression by microscopy. After 5-6 days the CHARMS or CHARMS-3xA transduced primary hematopoietic progenitor cells were FACS sorted for homogenous GFP expression levels.

### ELISA assay of IL-2 release from EL4 cells in response to IL-1β stimulation

To measure IL-2 release we used the Mouse IL-2 DuoSet ELISA kit (R&D Systems, DY402-05) using the manufacturer’s protocol. Briefly, 1x10^6^ cells in 150 μl per well were seeded into a 48-well plate. Cells were allowed to settle for 30 min, before being stimulated with recombinant mouse IL-1β (PeproTech) in 50 μl medium per well at a final concentration of 10 ng/ mL. For the unstimulated control 50 μl medium was added. After 24 hours plates were centrifuged (500x g for 5 minutes) and supernatants were transferred to a fresh 96 well plate. Supernatants were stored at -80°C until IL-2-ELISA analysis. Absorbance readings were acquired on a VersaMax Microplate Reader (Molecular Devices) at 450 nm. The obtained results were normalized based on the EL4-mScarlet-TRAF6/MyD88-GFP IL-2 release. IL-2 release was assayed in triplicates (n=3 technical replicates) on three independent days (N=3 independent biological replicates). IL-2 release was normalized to that of stimulated WT cells and baseline-corrected using unstimulated 3xKO. For data presentation and visualization the average of biological replicates is presented and shown (Fig. 1D, Fig. 2G, Fig 4F, fig. S1B and fig. S6C). For dose response of CHARMS-1x, 3x and 5x, the same stimulation and assay procedure was performed. CHARMS expressing 3xKO cells were stimulated with a IL-1β concentration range of 0.001 to 100 ng/ml (Fig. 4F). Here all the IL-2 values were normalized to the maximal and minimum IL-2 release (that is CHARMS-5x at 100 ng/ml of IL-1 and the lowest unstimulated CHARMS measurements).

### ELISA assay of IL-6 and TNF-ɑ release from primary bone marrow derived murine macrophages in response to TLR4, 7 and 9 stimulation

After seven days of differentiation, WT and 2xKO primary bone marrow derived murine macrophages were re-seeded into a TC-treated 96-well-plate at a density of 50x10^3^ cells / well. The next day cells were stimulated with the following TLR agonists: LPS (innaxon, 100ng/ml) was added for 4h to stimulate TLR4, Cpg-B (ODN1668, InvivoGen, 1µM) was added for 6h to stimulate TLR9. IL-6 release was measured in these cell supernatants. For stimulation of TLR7 Resiquimod (R848, Invivogen, 50ng/ml) was added for 8h to measure TNF-alpha release in the supernatants. Cell supernatants were analyzed using Mouse IL-6 ELISA MAX Deluxe Set or Mouse TNF-alpha ELISA MAX Deluxe Set (BioLegend). IL-6/TNF-alpha release was assayed in triplicates (technical replicates) on three independent days (independent experimental replicates). WT were included in all measurements as a positive control for TLR4, 7 and 9 stimulation. For data presentation and visualization the average of biological replicates (N=3) is presented and shown (Fig. 1E).

### Imaging Chambers and Supported Lipid Bilayers

SLBs were generated using methodology outlined in prior publications (*21*, *35*). SLB were composed of a phospholipid mixture consisting of 97.5% mol 1-palmitoyl-2-oleoyl-sn-glycero-3-phosphocholine (POPC), 2% mol 1,2-dioleoyl-sn-glycero-3-[(N-(5-amino-1-carboxypentyl)iminodiacetic acid)succinyl] (ammonium salt) (DGS-NTA), and 0.5% mol 1,2-dioleoyl-sn-glycero-3-phosphoethanolamine-N-[methoxy(polyethylene glycol)-5000] (PE-PEG5000). Stock solutions of lipids dissolved in chloroform were sourced from Avanti Polar Lipids and blended in glass round-bottom flasks. To remove chloroform, the glass round-bottom flask was subjected to rotary evaporation and then placed under vacuum for 2 hours The dried lipids were reconstituted in PBS, and small unilamellar vesicles (SUV) were created through multiple freeze-thaw cycles. Finally the lipid mixture was centrifuged at 35,000xg for 45 minutes. The resulting SUV suspension was stored at 4°C, and used for experiments for up to five days.

For live cell imaging, supported lipid bilayers were set up in 96-well glass bottom plates purchased from SWISS SCI (cat. #PS96B-G175). The plates underwent the following cleaning process. First, we washed glass wells with a 5% Hellmanex solution containing 10% isopropanol heated to 50°C for 30 minutes. This was followed by a 1-hour incubation with 5% Hellmanex solution at 50°C. Subsequently plates were washed extensively with pure MQ water, dried using nitrogen, and sealed with Polyolefin tape (Thermo Scientific). On the day of the experiments, individual wells in the 96-well plate were excised and then base-etched for 15 minutes with 5 M KOH, washed with water, and then equilibrated with PBS. SUVs suspensions were deposited in each PBS containing well and allowed to form for 1 hour on a hot plate set at 45°C. Wells were then thoroughly washed with PBS and incubated for 20 minutes with HEPES buffered saline (HBS: 20 mM HEPES, 135 mM NaCl, 4 mM KCl, 10 mM glucose, 1 mM CaCl2, 0.5 mM MgCl2) containing 5 mM NiCl2 to charge the DGS-NTA lipid with nickel. Post-nickel charging, SLBs underwent washing in HBS 0.1% BSA and a subsequent 30-minute incubation. After blocking, SLBs were functionalized by incubating with His-tagged IL-1β for 1 hour, before being washed extensively with HBS 0.1% BSA. The final labeling concentration of His10-IL-1β was 5 nM, which resulted in a measured surface density of 20 to 50 His10-Halo-IL-1β molecules per µm^2^.

### Protein Expression, Purification and Labeling

We functionalized supported membranes with an engineered variant of mouse IL-1β (in text referred to as His10-Halo-JF646-IL-1β). This IL-1 variant had been engineered to have a 10x His tag, and could therefore label membranes that contained DGS-NTA lipids. Full details on the design, expression and purification of this construct have been previously reported (*21*, *28*). Briefly, His10-Halo-IL-1β was expressed and purified as a spycatcher-spytag conjugation of two separate engineered proteins (IL-1β-Spytag and His10-Halo-Spycatcher). The conjugated protein was purified to homogeneity using Immobilized metal affinity chromatography, anion exchange and gel filtration chromatography steps. For microscopy detection and analysis the HaloTag of this protein was labeled with Janelia Fluor 646 (JF646)-HaloTag Ligand (a gift from Luke Lavis, Janelia Research Campus HHMI). Labeling was performed overnight at 4 C followed by gel filtration over a Superdex 200 column. For microscopy calibration of mScarlet single molecule intensity we used His10-mScarlet-IL-1β. For mEGFP single molecule intensity calibration, mEGFP was expressed and purified as previously described (see Deliz et al., 2021 (*21*)).

### TIRF-Microscopy data acquisition

Imaging of EL4 cells expressing different CHARMS-GFP constructs and TRAF6-mScarlet was performed on an inverted microscope (Nikon TiE, Tokyo, Japan) equipped with NIKON fiber launch TIRF illuminator. Illumination was controlled with a laser combiner using the 488, 561 and 640 nm laser lines. Fluorescence emission was collected through filters for GFP (525 ± 25 nm), RFP (595 ± 25 nm) and JF646 (700 ± 75 nm). All images were collected using a Nikon Plan Apo 100x 1.4 NA oil-immersion objective that projected onto a Photometrics 95B Prime sCMOS camera with 2x2 binning and a 1.5x magnifying lens (calculated pixel size of 0.147 µm). Image acquisition was performed using NIS-elements software. All live cell microscopy was performed at 37°C. The microscope stage temperature was maintained using an OKO Labs heated microscope enclosure.

### Imaging mScarlet-TRAF6 recruitment to CHARMS on IL-1β functionalized SLBs with TIRF-Microscopy

His10-Halo-JF646-IL-1β functionalized SLBs were set up as described above. To quantify the IL-1β density on SLBs before each experiment, wells were prepared that were functionalized with identical labeling protein concentration and time, but with different molar ratios of labeled to unlabelled His10-Halo-IL-1β. Before application of cells, SLBs were analyzed by TIRF microscopy to check formation, mobility and uniformity. Short time series were collected at wells containing a ratio of labeled to unlabelled His10-Halo-IL-1β, (e.g. <1 His10-Halo-JF646-IL-1β molecule per µm2) to calculate ligand densities on the SLB based upon direct single molecule counting. All experiments were performed at IL-1 SLB densities of 20 to 50 molecules per µm2.

Before each imaging experiment we acquired calibration images using recombinant mEGFP and mScarlet-i. To image single GFP/mScarlet-i fluorophores, recombinant purified mEGFP and mScarlet-i (*28*) was diluted in HBS and adsorbed to KOH cleaned glass. Single molecules of GFP or mScarlet-i were imaged using identical microscope acquisition settings to those used for cellular imaging. To image live cells, EL4 cells were pipetted onto supported lipids bilayers functionalized with His10-Halo-JF646-IL-1β. EL4 cells expressing different CHARMS-GFP constructs and TRAF6-mScarlet were sequentially illuminated for 60 ms with 488 nm and 300ms with 561 nm laser line at a frame interval of 4 s.

### FRAP data acquisition and data analysis

FRAP experiments were performed at the Advanced Medical BioImaging Core Facility at the Charité, on a NIKON TIRF microscope (Nikon Eclipse Ti-E) with the FRAPPA module. Fluorescent images were acquired with a Nikon Plan Apo 100x 1.4 NA oil-immersion objective and projected on a Photometric Prime 95B sCMOS camera. For FRAP analysis of CHARMS, we prepared SLBs as described above. Before the addition of cells to the imaging chamber, we analyzed SLB formation and mobility by visual inspection of the fluorescently labelled IL-1β. We prepared EL4 cells for imaging by washing in PBS and resuspending in HBS To ensure FRAP analysis of fully assembled signalosomes, we incubated cells with IL-1β functionalized SLBs for at least 30 mins before image acquisition. After incubation, we selected EL4-3xKO cells containing multiple GFP labeled CHARMS signalosomes for FRAP analysis. The photobleach spot was centered on large stationary signalosomes. Images were recorded 16 sec before and 120 sec after photobleaching at a time interval of 1 image per 4 seconds. All fluorescent images were acquired using an exposure time of 60-100 ms.

To quantify FRAP recovery we determined the mean intensity of the photobleached region as a function of time. The background intensity was measured from a cell free region and was subtracted from all timepoints. We calculated the rate of photobleaching by measuring the intensity in a region of interest centered on a stable signalosome. This was used to generate a bleaching correction curve that was used to correct the intensity of the bleached region. The data was normalized to the pre-bleach intensity using the following equation: Intensity(t)normalized=(Intensity(t)-Intensity(0))/(Intensity(pre-bleach)-Intensity(0)), where Intensity(pre-bleach) is the intensity preceding photobleaching, and Intensity(0) is the intensity immediately preceding photobleaching. Measurements from multiple photobleached signalosomes were averaged, and the standard deviation was calculated (Fig. 4E). We conducted three independent experimental replicates on different days for each FRAP experiment (e.g. for all CHARMS variants tested, Fig 4A-D).

### Analysis of RelA nuclear translocation

To assess whether CHARMS expression restored NF-kB activation, we analyzed the nuclear translocation of NF-kB subunit RelA in response to IL-1β stimulation. First, EL4-3xKO-mScarlet-TRAF6 cells expressing CHARMS variants were plated on poly-L-lysine coated 96 well plates and allowed to adhere for 30 mins. Cells were then stimulated with soluble IL-1β at a concentration of 10 ng/ml for 30 min. Cells were then fixed with 3.5% (wt/vol) PFA containing 0.5% (wt/vol) Triton X-100 for 20 min at room temperature. Cells were washed with PBS and blocked with PBS 10% BSA (wt/vol) at 4°C overnight. The next day, fixed cells were labeled with staining solution consisting of anti-RelA (1:400, Cell Signaling Technology, #8242) and Nano-secondary AF647 (Chromotek, #srbAF647-1, 1:1000) diluted in PBS 10% (wt/vol) BSA containing 0.1% (vol/vol) Triton X-100. Cells were incubated for 1 h at room temperature. Then, cells were counterstained with solution containing RelA (1:300), Nano-secondary AF647 (1:750), Hoescht (to label nucleus) and 488-phalloidin (to label the cell contour) for 30 min at room temperature. Finally, cells were washed with PBS. After labeling, wells containing stained cells were maintained in PBS.

We acquired spinning disk confocal images of RelA nuclear translocation at the Advanced Medical BioImaging Core Facility at the Charité. Spinning disk confocal was performed on an inverted microscope (Nikon TiE) equipped with the appropriate laser lines (e.g., 405, 488 and 638 nm laser lines). Fluorescent images were acquired with Nikon Plan Apo 60x 1.4 NA oil objective lens and projected onto a Hamamatsu ORCA-Fusion digital sCMOS camera (calculated pixel size of 107 nm). Image acquisition was performed with NIS-Elements software.

We quantified spinning disk confocal images of RelA nuclear localization in an analysis pipeline implemented in FIJI and Cell Profiler, previously reported here (*28*). Briefly, first we performed background subtraction from the GFP and RelA (Cy5 channel) immunofluorescence staining micrographs in FIJI. Background was removed in two steps: First, we subtracted a dark field image from each image. Secondly we estimate cytosolic background by generating median blur from each micrograph. We then subtracted this median blur from the parent micrograph. We then performed segmentation and quantification using a Cell Profiler pipeline. This pipeline first segmented the cell nucleus using the Hoechst channel. Segmented nuclei had to have a diameter between 50 to 130 pixels to be retained for further quantification This filtering step was necessary to exclude small Hoescht stained objects that corresponded to cell fragments and apoptotic cells. Next we segmented the GFP/Phalloidin channel and identified the total cell volume. These segmentation steps were performed using an Otsu threshold. The volume corresponding to cellular cytoplasm is identified by subtracted the total cell volume minus nucleus volume. The RelA staining intensity of the cell nucleus was normalized to the RelA cytoplasm intensity (e.g. RelA nuclear intensity divided by the RelA cytoplasmic intensity). Finally, we performed data visualization of the normalized RelA nucleus intensity in ggplot2 (Fig. 1C).

### AlphaFold2-Complex structure predictions of *Nostoc sp. 106C* bacterial Death Domain

We used the AlphaFold2Complex (AF2C) software (*31*) to predict oligomeric structural models of the bacterial Death Domain from *Nostoc sp. 106C* (Accession number WP_086756477.1). First, to predict the domain boundaries, we created a multiple sequence alignment of the FASTA file of the previously annotated bDD found in CHAT domain-containing proteins (*8*). Based on this alignment we used WP_086756477.1[aa230-303] as the input sequence to AlphaFold2-Complex and to generate the CHARMS-bDD (Fig. 2D, fig. S2A,B). We collected input features and conducted a multiple sequence alignment to prepare the oligomeric predictions using AF2C. We used the preset Super2 for all oligomeric predictions. Different oligomeric stoichiometries are fed to five neural networks each (*AlphaFold Multimer* v3) and five models are predicted for each oligomer setup. AlphaFold Multimer predictions were run on the Max Planck High Performance Computing Centre. The final *pdb* files and the output summary *json* files are saved. We assessed the helical filament formation visually and used the iScore and piTM metric, two outputs of the AF2C algorithm, to assess the confidence of oligomeric models in complex formation.

### Visualization of protein filament structure

We generated symmetrical helical filament models for the MyD88-DD, *Nostoc sp. 106C* bDD (WP_086756477.1) and DHF91 (*23*) using the Rosetta macromolecule modeling suite (Rosetta 3.13). For the visualization we generated all symmetrical helical models with comparable length (∼130 Å). We generated the MyD88-DD helical filament model using Cryo-EM structure (PDB-ID: 6I3N, (*15*)) as a starting reference (Fig. 2A). Due to no experimental derived structure, we created a starting reference of bDD by generating a decameric oligomer model in AlphaFold2-Complex (*31*). We used the AFC2 predicted decameric oligomeric model with highest iScore and piTM metrics to create the helical filament visualization (Fig. 2D).

To create the symmetrical helical filaments shown (Fig 2), we relaxed these experimentally or computationally derived structural models using the Rosetta energy function. We added a coordinate constraint to the backbone heavy atoms of these initial structures (using the *-relax:constrain_relax_to_start_coords* operation in the command line). We selected the relaxed oligomeric models with the lowest scored energy. We then selected two neighboring subunits along the helix from these relaxed structures to build pdb files of open-ended symmetrical helical filament using the *make_symmdef_file.pl* function in Rosetta (*36*). The DHF91 structure (PDB-ID: 6E9X, (*23*)) was sufficiently long and therefore did not need to be further extended (Fig. 2E).

The generated pdb files for these helical filaments were visualized using ChimeraX. The structures are displayed in the ribbon format and colored with the same color scheme used for the schematics and data plots of CHARMS construct composed of these domains. The MyD88-DD filament shown in Figure 2B contains 21 subunits (129 Å), the bDD filament 23 subunits (131 Å) and the DHF91 filament 15 subunits (128 Å). The outer diameter and the length of the filaments was determined in ChimeraX using the “Distances” tool.

To visualize the confidence of the AlphaFold2 generated bDD model, one subunit of the bDD monomer in the decameric helical filament was rendered with a LUT that showed the per-residue confidence metric (predicted Local Distance Difference Test - pLDDT values). We aligned the bDD monomer structural monomer with the MyD88-DD structures using the “Matchmaker” tool in ChimeraX. From this alignment the Root Mean Square Deviation (RMSD) for the C-alpha-atoms (RMSD of backbone) was calculated (fig. S2A-B). All structural figures were generated in ChimeraX.

### Quantification and Statistical Analysis

All data are expressed as the mean ± the standard deviation (SD) or mean ± the standard error of the mean (s.e.m.), as stated in the figure legends, captions and results. The exact value of n and what n represents is stated in figure legends and results. Means were compared using the unpaired t-test or one-way ANOVA with Tukey post-hoc test. Before using one-way ANOVA, data was checked for normality and equal variance using Shapiro-Wilk test and Levene’s test. The analysis scripts used in this study available at github.

### Image analysis and particle tracking of two color TIRF data

To quantify the dynamics of Myddosomes/CHARMS signalosomes and TRAF6-mScalet in live EL4 cells, we created an image analysis pipeline that runs in Fiji, Python and R. Image analysis scripts were written to run on cluster computers at the Max Planck Computing and Data Facility (Garching, Bavaria, Germany). The aim of this image analysis pipeline was to first process the images to remove intensity count that derived from camera noise or background fluorescence (non-specific cytosolic fluorescent signal), and then identify and track the individual fluorescently labeled signalosomes complexes within the live cell data.

To process the image stack we first subtracted a dark frame image (e.g. an image only containing intensity values from current and noise generated by the camera electronics) from each image file. The dark frame image was created by averaging 100 images acquired without light exposure to the camera and with identical acquisition settings to those used for live cells imaging. From this point dark frame subtracted images were processed to create two sets of image stacks: one set was used for intensity measurements to quantify the dynamics of signalosome assembly (referred to as the intensity reference image). The second set was processed to enhance the contrast of the punctate structure and was used to optimally identify and track signalosomes (referred to as the tracking images). To generate an images stack for tracking signalosome puncta, the GFP/mScarlet images were processed with a median filter with a 11-pixel diameter. The resulting median filtered image is subtracted from the original to remove background intensity. We next applied two methods to remove stochastic fluctuations associated with the fluorophore blinking or camera read noise, both of which can contribute to errors in particle detection and linking detected particles between frames. First, we applied a second median blur of 5-pixel diameter to even out the field. Because median filters are edge-preserving (keeps silhouette), a 5-pixel median-blur preserved punctate structures while further removing background noise. We then applied a moving average of 3 frames to further suppress intensity fluctuations and smooth the image. Because we were interested in detecting signalosomes (e.g. puncta that possibly contain both Myddosomes/CHARMS and TRAF6) we reasoned that combining the mGFP and mScarlet channels into a single image would further enhance the detection and tracking of signalosome fluorescent puncta. Therefore all mGFP and mScalet-i image frames were added together. The result was a single time series with high signal to noise and optimized for particle detection and tracking.

The combination of two median blur filtering steps and running average used to create the tracking reference image skewed intensity values. Therefore to be able to accurately quantify signalosome intensity and assembly kinetics we generated an intensity reference image that preserved intensity values. To accomplish this, we estimate cytosolic background using a median blur with a diameter of 25-pixels, which is roughly half the diameter of the cell:SLB interface observed in the TIRF field. This median blurred image represents an estimate of background intensity from the cytoplasmic fluorescence (that is CHARMS-GFP/TRAF6-mScarlet not assembled into signalosomes). This estimated background fluorescent intensity is subtracted from the dark-frame removed images and the resulting image is the intensity reference image from which all analyzed intensity values are derived.

Next, individual cells within the time series were identified using a marker-controlled watershed segmentation implemented on a maximum projection image of the GFP intensity channel. The centroids of the segmented cells were used to break both the tracking image and intensity reference image series into small image series of individual cells. We used the Fiji TrackMate plugin to track the signalosome puncta within the tracking image for each segmented cell. We tracked signalosome puncta with a maximum linkage distance of 5 pixels, a tracking threshold of 2 and a tolerance of 2 gap-closing frames between detected particles. After processing in Trackmate, tracking coordinates generated were imported into *R*. To fill in missing frames (e.g. gap-closing frames with no detected particles) within tracks, we approximated the location of the missing puncta using the Euclidean distance from the coordinate of the preceding and following detected particle. We extracted the fluorescence intensity of puncta from the intensity reference image for both the MyD88-GFP/CHARMS-GFP image and the TRAF6-mScarlet image. The intensity was measured from a 5x5 circular pixel region center of the puncta centroid. To achieve sub-pixel accuracy, we segmented a square matrix of pixels from images. This square was centered on the tracked puncta centroid. The segmented area was expanded 11-fold with the original pixel intensity subdivided in the expanded area pixels. The square was then overlaid with a circle mask (diameter of 5 pixels that was also expanded by 11-fold). The pixel intensities overlaid by this circle mask in the GFP and mScarlet channel were summed to give a puncta intensity for both MyD88-GFP/CHARMS-GFP or mScarlet-TRAF6 channel respectively.

### Quantification and analysis of CHARMS dynamics and TRAF6 recruitment

To estimate the number of CHARMS monomers and TRAF6 within tracked signalosomes, the fluorescent intensity was divided by the median intensity of single mEGFP or mScarlet-I fluorophores respectively, using a methodology previously described (*28*, *29*). Briefly, to compute the distribution of single GFP/mScarlet-i fluorophores intensities, we processed images of single mGFP or mScarlet fluorophores absorbed to glass identically to CHARMS and mScarlet-TRAF6 images. To measure the maximum normalized intensity and use this as a metric for signalosome size or the multimerization of TRAF6, we limited the analysis to the first 100 frames of tracked CHARMS-GFP and mScarlet-TRAF6 puncta. Additionally, we only included the first 200 frames, after the cell had landed on the bilayer, in the analysis. These restrictions were chosen to limit the effect of photobleaching on the quantification of signalosome sizes.

Fluorescent GFP puncta that represented CHARMS signalosomes were scored positive for TRAF6 recruitment when the mScarlet-TRAF6 normalized fluorescence intensity was ≥ 1.5. To assay size and extent of TRAF6 multimerization at CHARMS signalosome, we used the TRAF6 maximum intensity that had been normalized to the mean intensity of single mScarlet fluorophores. From this we plotted the distribution of TRAF6 multimers size at CHARMS signalosomes (Fig. 4C, and fig. S5A).

The lifetime of TRAF6 recruitment to CHARMS signalosomes was calculated by taking the sum of consecutive frames where the TRAF6 signal was above the fluorescent intensity threshold (≥ 1.5 of the normalized fluorescence intensity, Fig. 4D). Long-lived TRAF6 recruitment events were defined as a lifetime ≥40 s. The distribution of the percentage of long recruitment events to CHARMS signalosomes per cell is plotted in Fig 2F and Fig 4D. Distributions of recruitment events lasting ≤4s, 4-40s and ≥40s are provided in the supplemental figures (fig S2C-D, fig S4C-D). The same ranges were used to calculate the proportion of all observed recruitment events and plotted in Fig S2C and S5C using a bar graph.

To compare the probability of TRAF6 recruitment across different CHARMS signalosomes and examine the relationship between signalosome size and TRAF6 recruitment, we computed the cumulative density function (CDF) for the frequency of TRAF6 recruitment versus CHARMS signalosome size. This calculation was carried out for each replicate separately. The mean of the replicates ± the s.e.m. is plotted in Fig. 4E and S3. We performed all data visualization described above using ggplot2, a data visualization package for *R* and Graphpad.

**Table S1:**
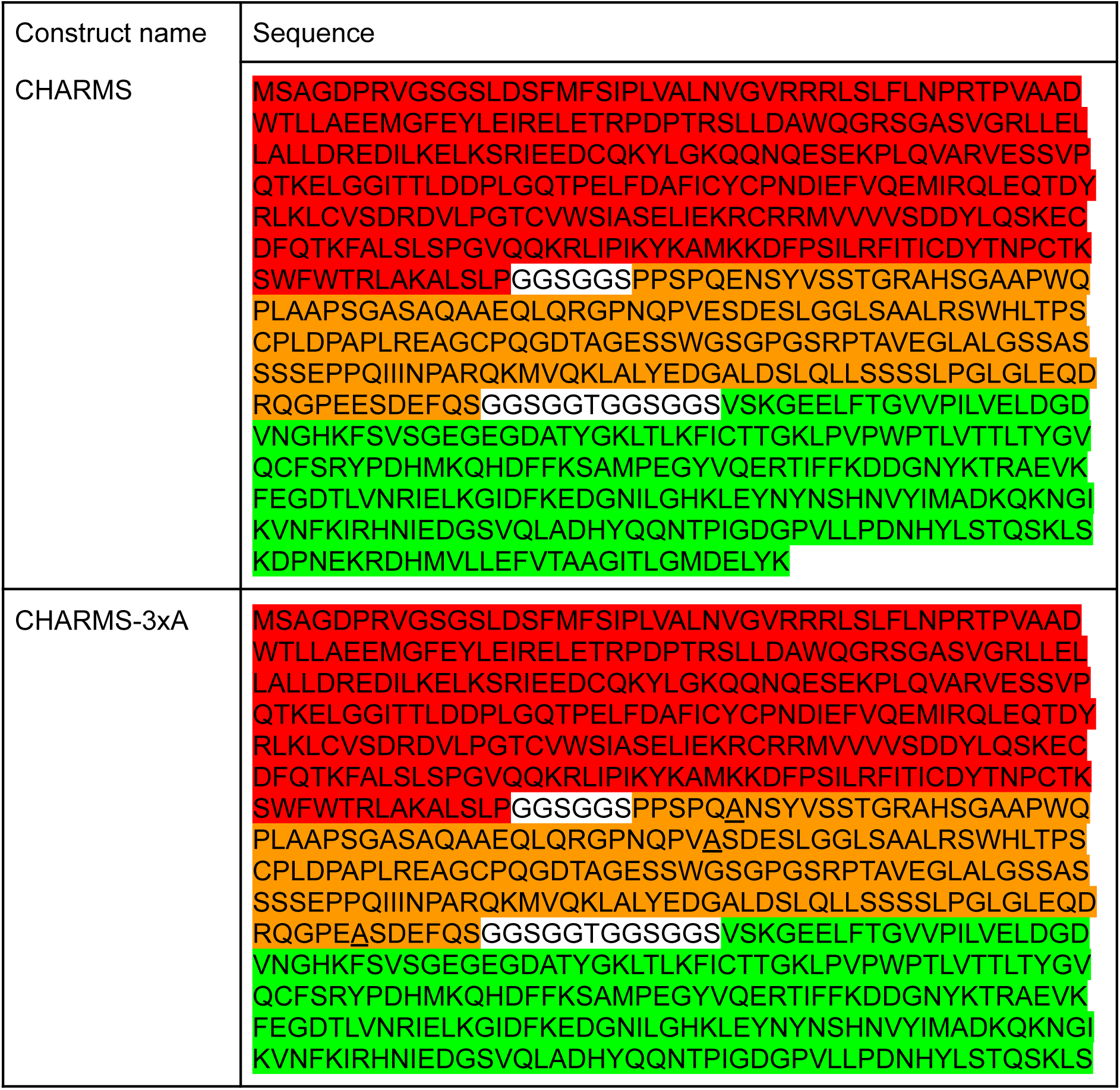
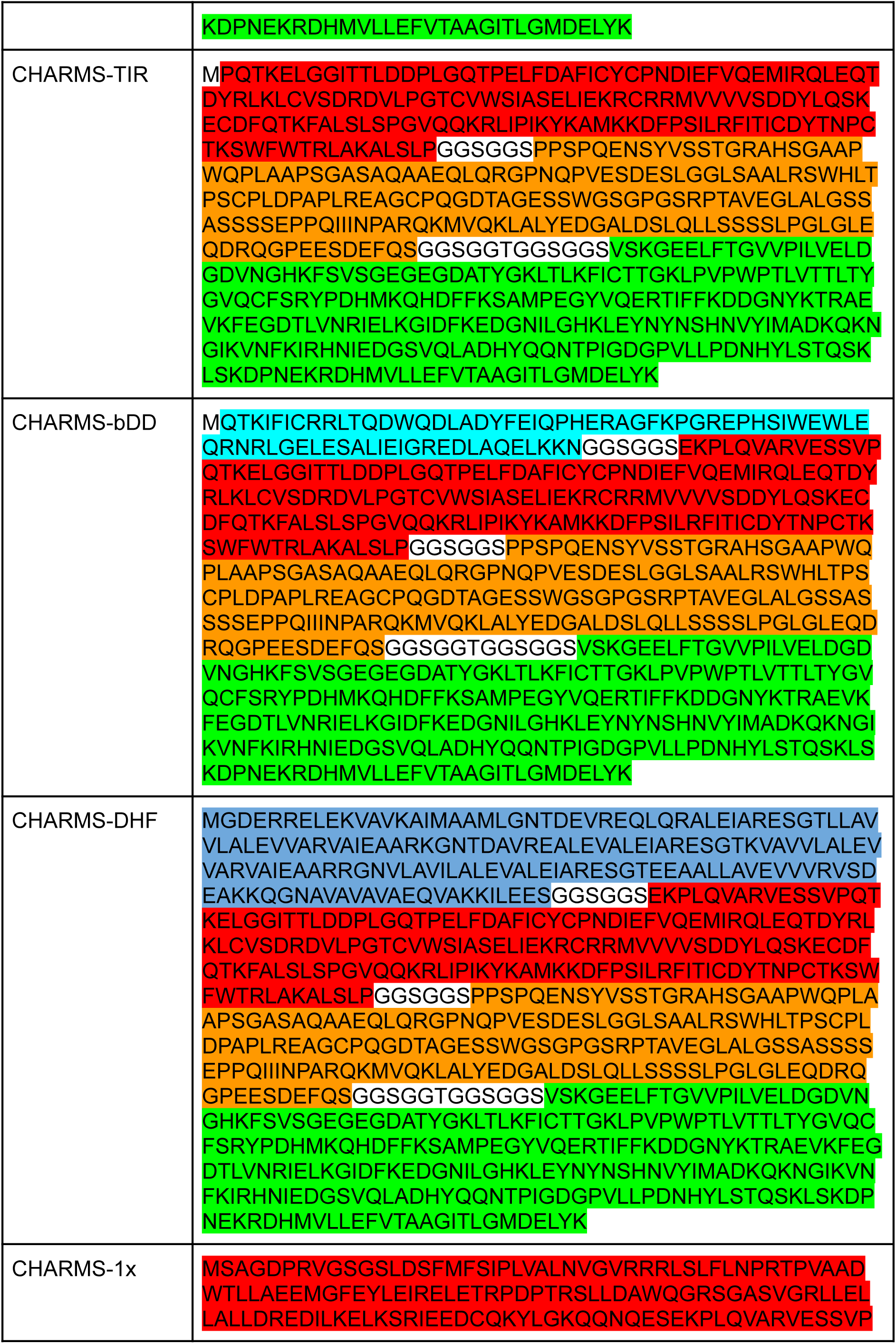
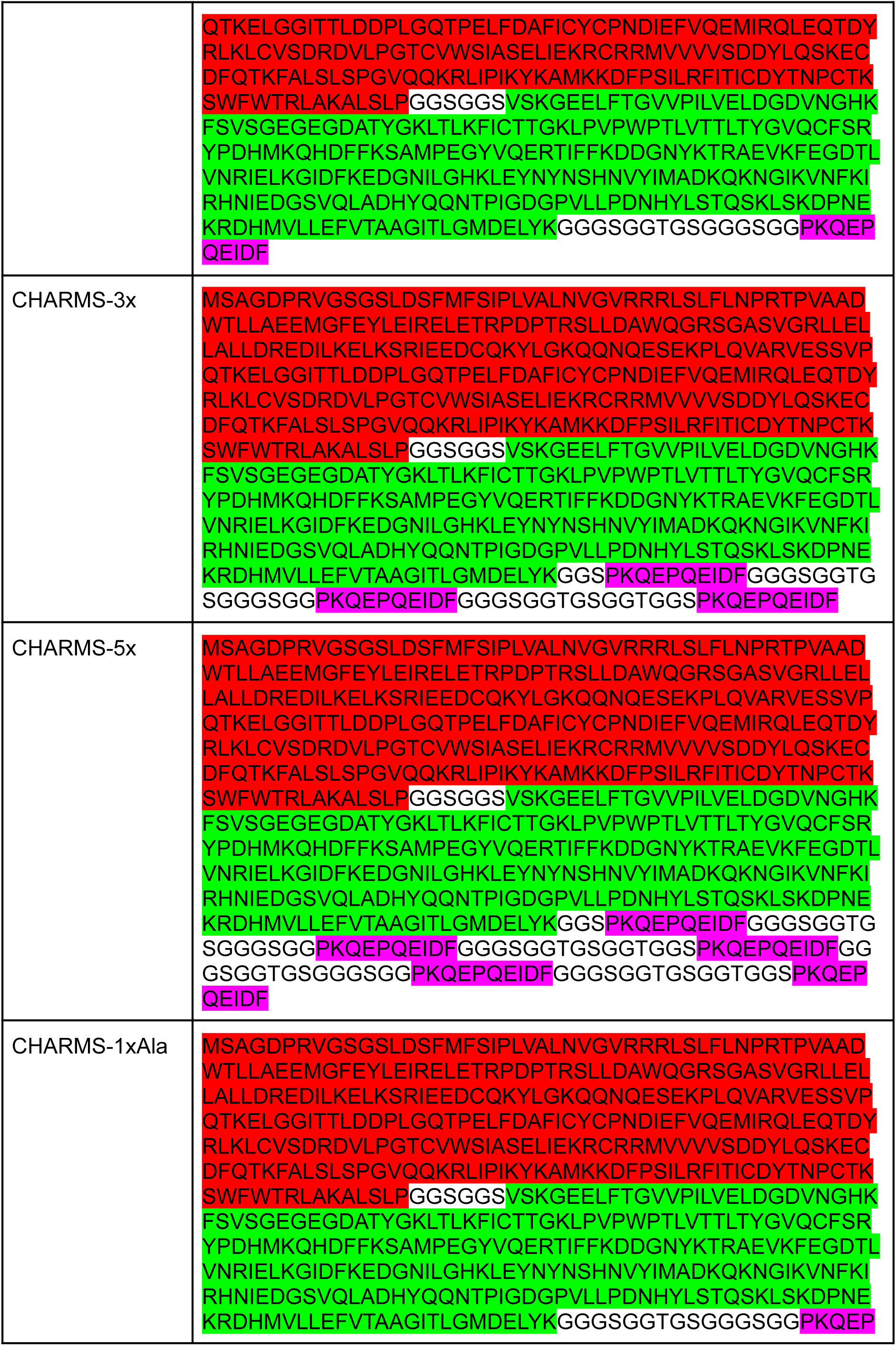
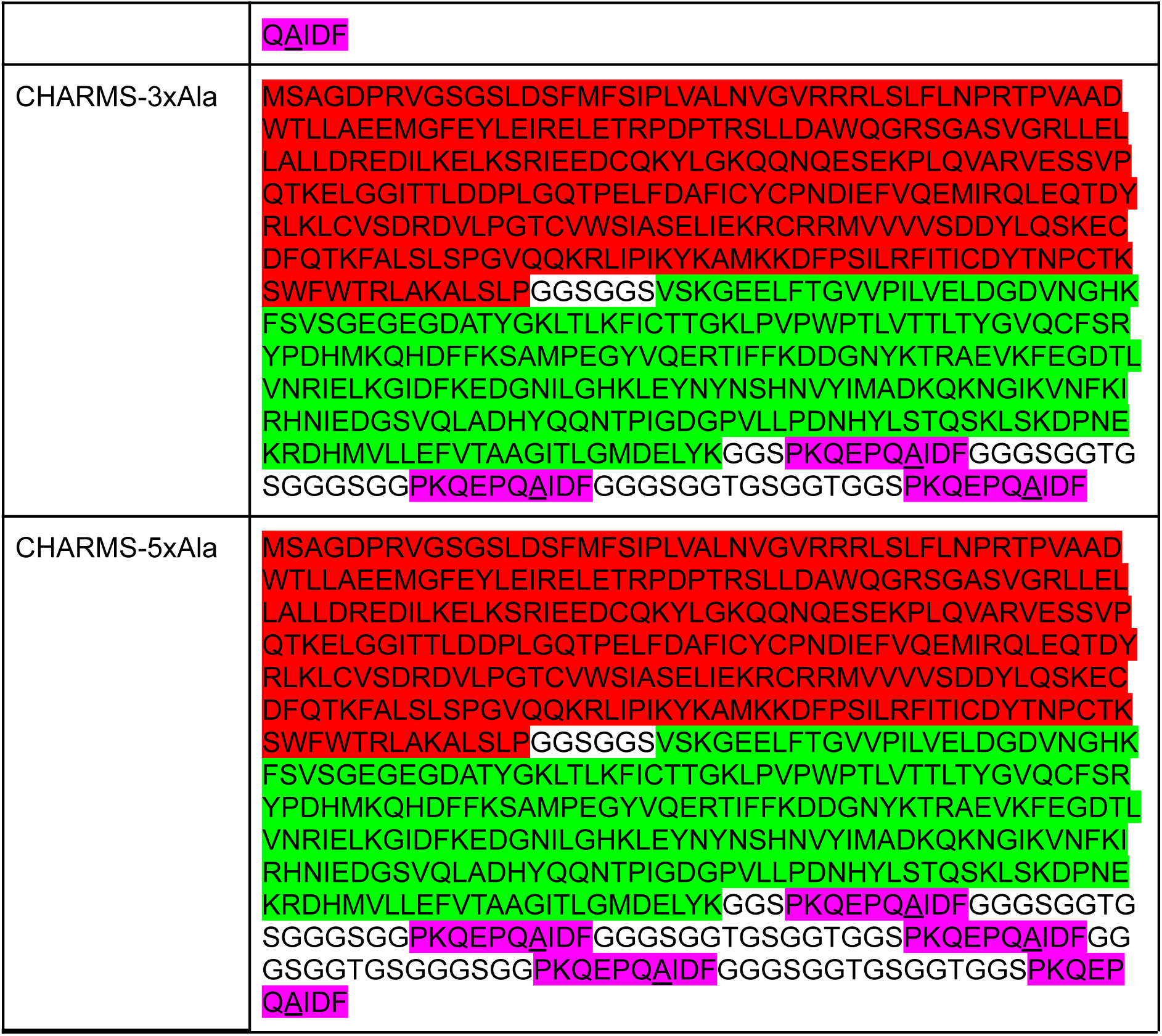
Complete amino-acid sequences and details of chimeric protein sequences of the CHARMS variants used in this study. Key: MyD88, IRAK1, mEGFP, bDD, DHF, TRAF6 binding motif (from CD40). <u>Point mutations are underlined</u>

**Table S2:** Identifiers for the origin of all elements used in this study.

| Element name | Identifier |
| --- | --- |
| MyD88 | ENSMUSG00000032508 |
| IRAK1<sup>538-712</sup> | ENSGT00940000160502 |
| DHF91 | Uniprot ID: 6E9X |
| bDD<sup>230-303</sup> | WP_086756477.1 |
| CD40<sup>229-238</sup> | ENSGT00940000161464 |

## Supplementary Movies

**Movie 1. Recruitment of E3 ligase TRAF6 to Myddosomes in WT cells**

Typical examples of Myddosome formation (labeled with MyD88-GFP) and TRAF6 recruitment. This movie shows an EL4 cell endogenously expressing MyD88-GFP and mScarlet-TRAF6 interacting with IL-1β functionalized SLBs imaged with TIRF microscopy. Same cell as shown in Fig 1F. Scale bar, 2 µm.

**Movie 2. Recruitment of E3 ligase TRAF6 to CHARMS signalosomes in 3xKO cells** Typical examples of CHARMS signalosomes and TRAF6 recruitment This movie shows a 3xKO cell endogenously expressing mScarlet-TRAF6 and reconstituted with CHARMS interacting with IL-1β functionalized SLBs image with TIRF microscopy. Same cell as shown in Fig 1G. Scale bar, 2 µm.

**Movie 3. CHARMS-TIR assembles in response to IL-1β stimulation but only transiently recruits TRAF6.**

This movie shows a 3xKO cell endogenously expressing mScarlet-TRAF6 and reconstituted with CHARMS-TIR interacting with IL-1β functionalized SLBs image with TIRF microscopy. CHARMS-TIR forms assemblies at the plasma membrane, but only recruits TRAF6 transiently. Same cell as shown in Fig 2B. Scale bar, 2 µm.

**Movie 4. CHARMS-bDD form signalosomes that recruit TRAF6 in response to IL-1β stimulation.**

This movie shows a 3xKO cell endogenously expressing mScarlet-TRAF6 and reconstituted with CHARMS-bDD interacting with IL-1β functionalized SLBs image with TIRF microscopy. CHARMS-bDD forms signalosomes at the plasma membrane that stably recruit TRAF6. Same cell as shown in Fig 2C. Scale bar, 2 µm.

**Movie 5. CHARMS-DHF form signalosomes that recruit TRAF6 in response to IL-1β stimulation.**

This movie shows a 3xKO cell endogenously expressing mScarlet-TRAF6 and reconstituted with CHARMS-DHF interacting with IL-1β functionalized SLBs image with TIRF microscopy. CHARMS-DD forms signalosomes at the plasma membrane that stably recruit TRAF6. Same cell as shown in Fig 2C. Scale bar, 2 µm.

**Movie 6. CHARMS with 1x, 3x, 5x TRAF6 binding motifs assemble into signalosomes that recruit TRAF6 in response to IL-1β stimulation.**

This movie shows a 3xKO cell endogenously expressing mScarlet-TRAF6 and reconstituted with CHARMS-1x (top row), CHARMS-3x (middle row) or CHARMS-5x (bottom row) interacting with IL-1β functionalized SLBs image with TIRF microscopy. Regardless of the multiplicity of T6BM, CHARMS-1x, 3x and 5x form signalosomes at the plasma membrane that recruit TRAF6. Same cells as shown in Fig 4b. Scale bar, 2 µm.

**Figure S1.**
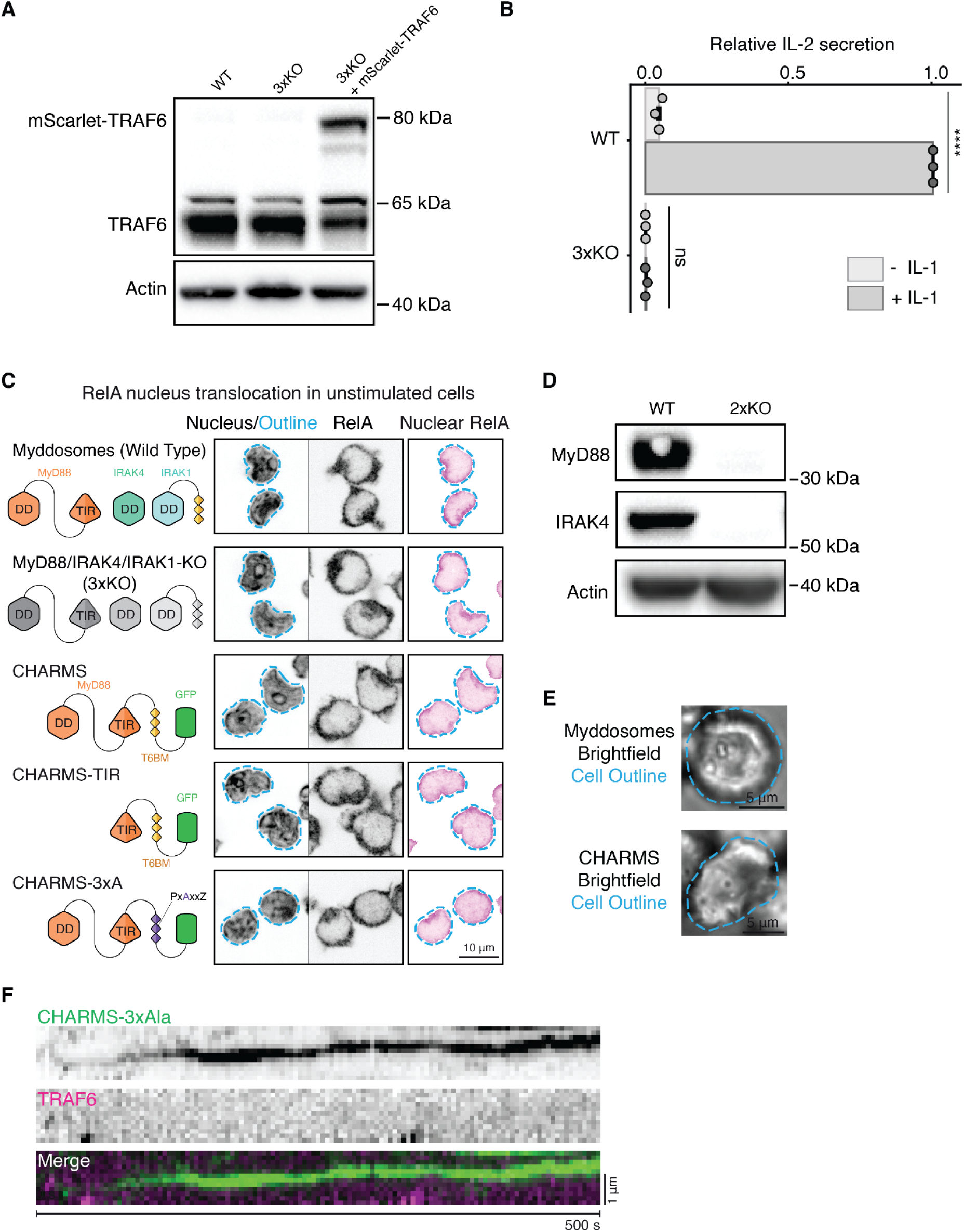
Cell line engineering to test CHARMS variants. **(A)** Western blot of CRISPR/Cas9 gene edited EL4-3xKO cells showing the heterozygous tagging of TRAF6 with mScarlet. **(B)** IL-2 ELISA showing that the EL4-3xKO-mScarlet-TRAF6 cells are unable to signal in response to IL-1. The IL-2 signal was normalized to the signal of the IL-1 stimulated EL4 WT cells. **(C)** Confocal micrographs showing the localization of RelA without IL-1 stimulation in WT, 3xKO cells, and 3xKO cells expressing CHARMSs variants. Dashed cyan outline superimposed on the Hoechst stain nucleus images shows segmented nuclear outline. This was used to segment and measure the RelA nuclear staining (third column). The quantification of this data is shown in Fig. 1C. Scale bar, 10 µm. **(D)** Western Blot showing the successful knockout of MyD88 and IRAK4 in bone marrow derived macrophages from MyD88/IRAK4 KO mice. The EL4 WT cell line is used as a positive control and actin as a loading control. **(E)** Brightfield images of the cells shown in Fig. 1E (top) and Fig. 1F (bottom). Brightfield images used to segmented the cell contours which are shown as dashed cyan lines superimposed on images in Fig. 1E, F. Scale bar, 5 µm. **(F)** IL-1 stimulation triggers the assembly of CHARMS-3xAla into plasma membrane puncta that do not recruit TRAF6. Kymograph analysis shows an example of CHARMS-3xAla assembly without TRAF6 recruitment. Scale bar, 1 µm.

**Figure S2.**
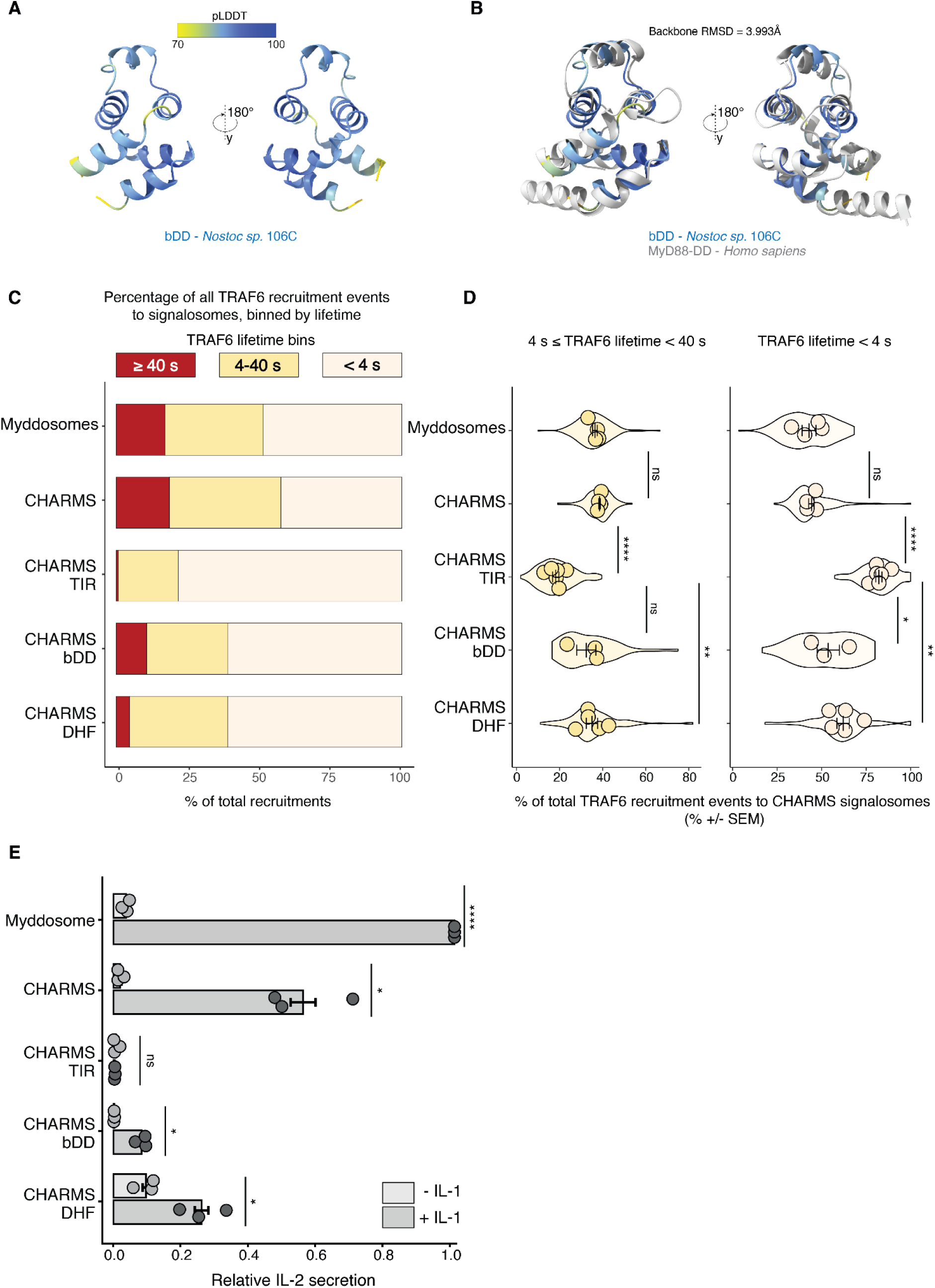
Bacterial Death-like Domain and computationally designed polymer forming proteins can functionally rescue stable TRAF6 recruitment. **(A-B)** The bacterial Death like Domains from Nostoc sp. 106C is predicted to fold into the typical 6 helix DD-superfamily fold (*5*). AlphaFold predictions of bacterial Death like Domains from *Nostoc sp.* 106C display high confidence (A). The predicted structure shows close structural similarities with the cryo-EM structure of human MyD88 Death Domain (PDB-ID: 6I3N) as assayed by the backbone RMSD of both structures (B). **(C)** Quantification of TRAF6 lifetime at Myddosomes and CHARMS signalosomes. TRAF6 recruitment was binned by lifetime into the following groups ≥40 s, 4-40 s and <4 s. The percentage of total for each TRAF6 lifetime bin is shown. **(D)** Quantification of TRAF6 lifetime at CHARMS signalosomes for recruitment events with<4 s (left) or 4-40 s (right) lifetimes. Plots show the percentage of the total TRAF6 recruitment events for each lifetime bin. Violin plots show the distribution of the percentage per individual cell across replicates. Mean percentage of TRAF6 lifetime <4 s ± s.e.m.: 43.0±3.9%, 44.3±1.5%, 82.2±1.7%, 53.7±6.3%, 62.1±3.5% for WT, CHARMS, CHARMS-TIR, CHARMS-bDD and CHARMS-DHF respectively. Mean percentage of TRAF6 lifetime 4-40 s ± s.e.m.: 36.4±1,1%, 38.6±0.4%, 18.2±1.4%, 32.4±4.5%, 35.0±2.6% for WT, CHARMS, CHARMS-TIR, CHARMS-bDD and CHARMS-DHF respectively. Data points superimposed on the violin plots show the mean of individual experimental replicates, n = 4, 4, 6, 3, and 5 replicates for cells expressing Myddosomes, CHARMS, CHARMS-TIR, CHARMS-bDD and CHARMS-DHF. The P-value are * p < 0.05; ** p < 0.01. Statistical significance is determined using unpaired t-test. **(E)** ELISA measurement of IL-2 release in EL4-MyD88-GFP/mScarlet-TRAF6 cells expressing all Myddosome components (referred to as WT in the figure) and 3xKO cells expressing CHARMS variants. Here the same data as shown in Fig 2G is normalized to the IL-2 release of the stimulated WT cells. WT, CHARMS and CHARMS-TIR ELISA data the same as presented Fig. 1C, and shown here for comparison to CHARMS-bDD and CHARMS-DHF. The P-value are **** =0.0001 and ** =0.01. Bars represent mean ± s.e.m (mean calculated from n=3 independent experimental replicates). Statistical significance is determined using unpaired t-test.

**Figure S3.**
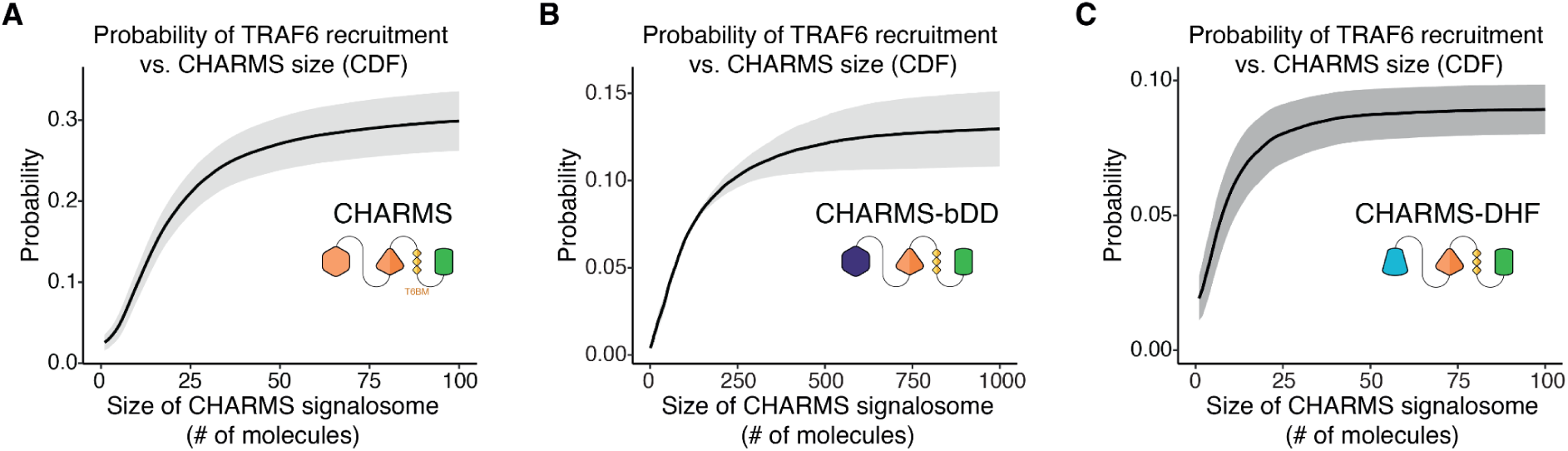
Probability of TRAF6 recruitment increases with size of CHARMS signalosomes. **(A-C)** The cumulative distribution function (CDF) of probability of TRAF6 recruitment versus the size of CHARMS signalosomes in CHARMS (A), CHARMS-bDD (B) and CHARMS-DHF (C). Colored lines show the mean of three independent experimental replicates (n = 3-5) and shaded area indicates S.E.M.

**Figure S4.**
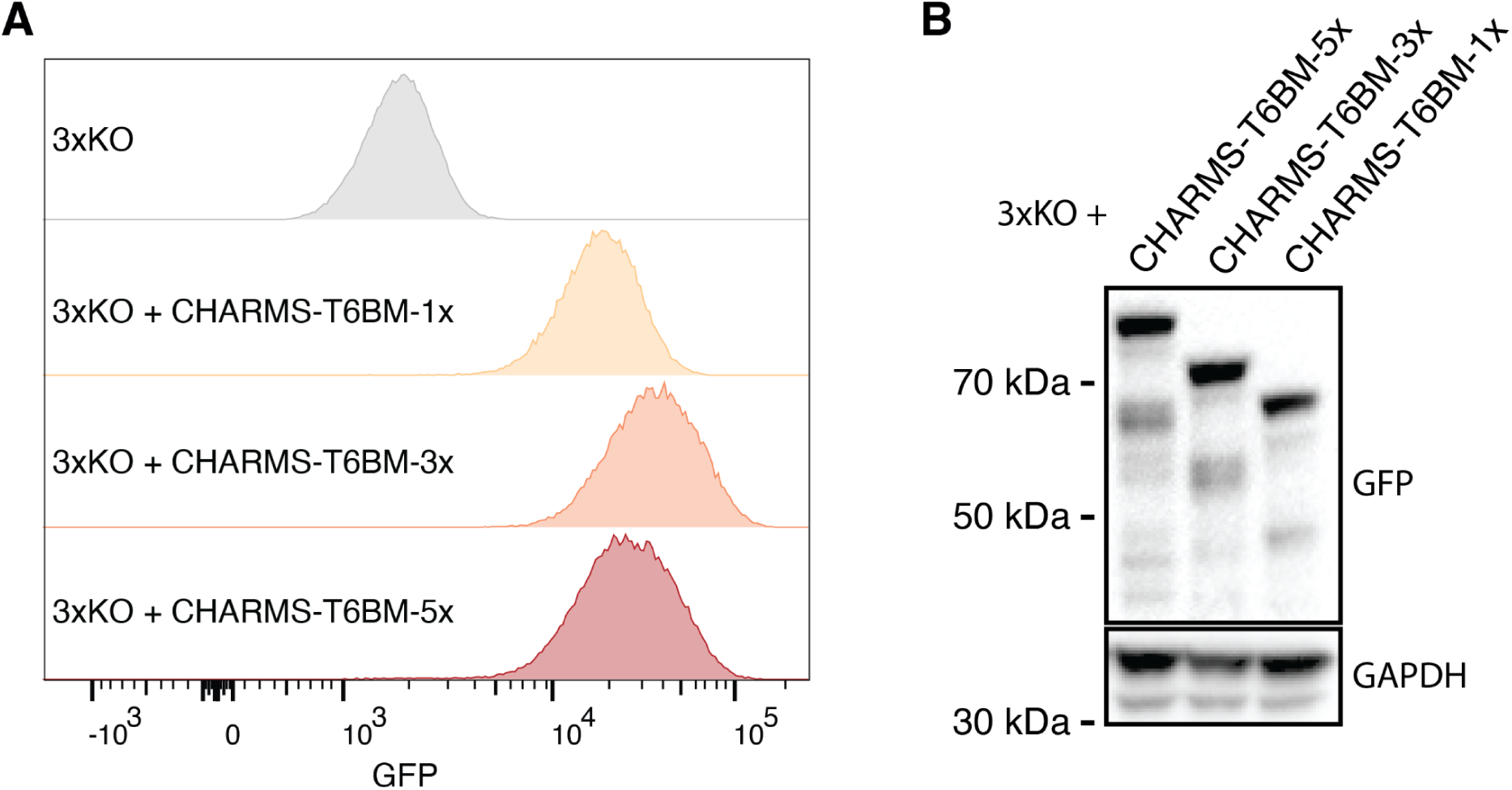
Engineering of cell lines with varied TRAF6 binding motif number. **(A)** Flow cytometry data comparing the expression level of the CHARMS-1x, -3x and 5x in the 3xKO cell line. **(B)** Western Blot confirming the similar expression level of the CHARMS constructs in the 3xKO cell line.

**Figure S5.**
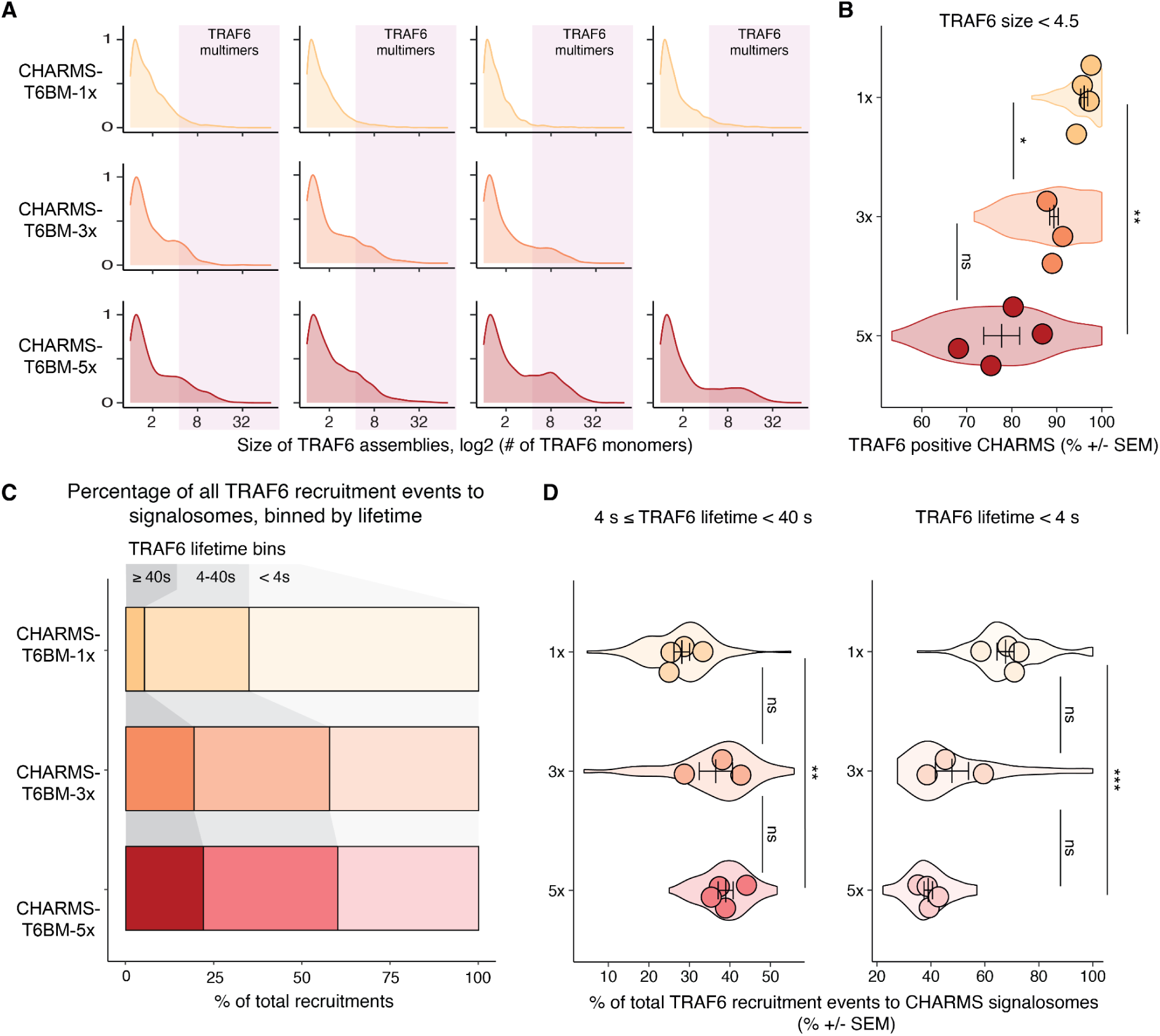
TRAF6 recruitment to CHARMS signalosomes is modulated by the multiplicity of the TRAF6 binding motif. **(A)** Density plots showing the distribution of TRAF6 assembly sizes that form at CHARMS signalosomes. Each density plot from an independent experimental replicate. Light magenta shaded area represents TRAF6 assemblies classified as multimers (that is containing ≥ 4.5 TRAF6 molecules). **(B)** Quantification of the percentage of CHARMS signalosomes associated with TRAF6 assemblies composed of <4.5 TRAF6 molecules. Violin plots show the distribution of individual cell measurements. Colored dots superimposed on violin plots correspond to the average value per cell across independent experimental replicates. P values, * p < 0.05; ** p < 0.01. Quantifications are from n = 3-4 replicates with 11-29 cells measured per replicate. Bars represent mean ± S.E.M. Statistical significance is determined using unpaired t-test. **(C)** Quantification of TRAF6 lifetime at CHARMS signalosomes. TRAF6 recruitment was binned by lifetime into following groups: ≥40 s, 4-40 s and <4 s. The percentage of total for each TRAF6 lifetime bin is shown. **(D)** Quantification of TRAF6 lifetime at CHARMS signalosomes for recruitment events with<4 s (left) or 4-40 s (right) lifetimes. Plots show the percentage of the total TRAF6 recruitment events for each lifetime bin. Violin plots show the distribution of individual cell measurements. Colored dots superimposed on violin plots correspond to the average value per cell across independent experimental replicates. P values, * p < 0.05; *** p < 0.001. Quantifications are from n = 3-4 replicates with 12-36 cells measured per replicate. Bars represent mean ± S.E.M. Statistical significance is determined using unpaired t-test.

**Figure S6.**
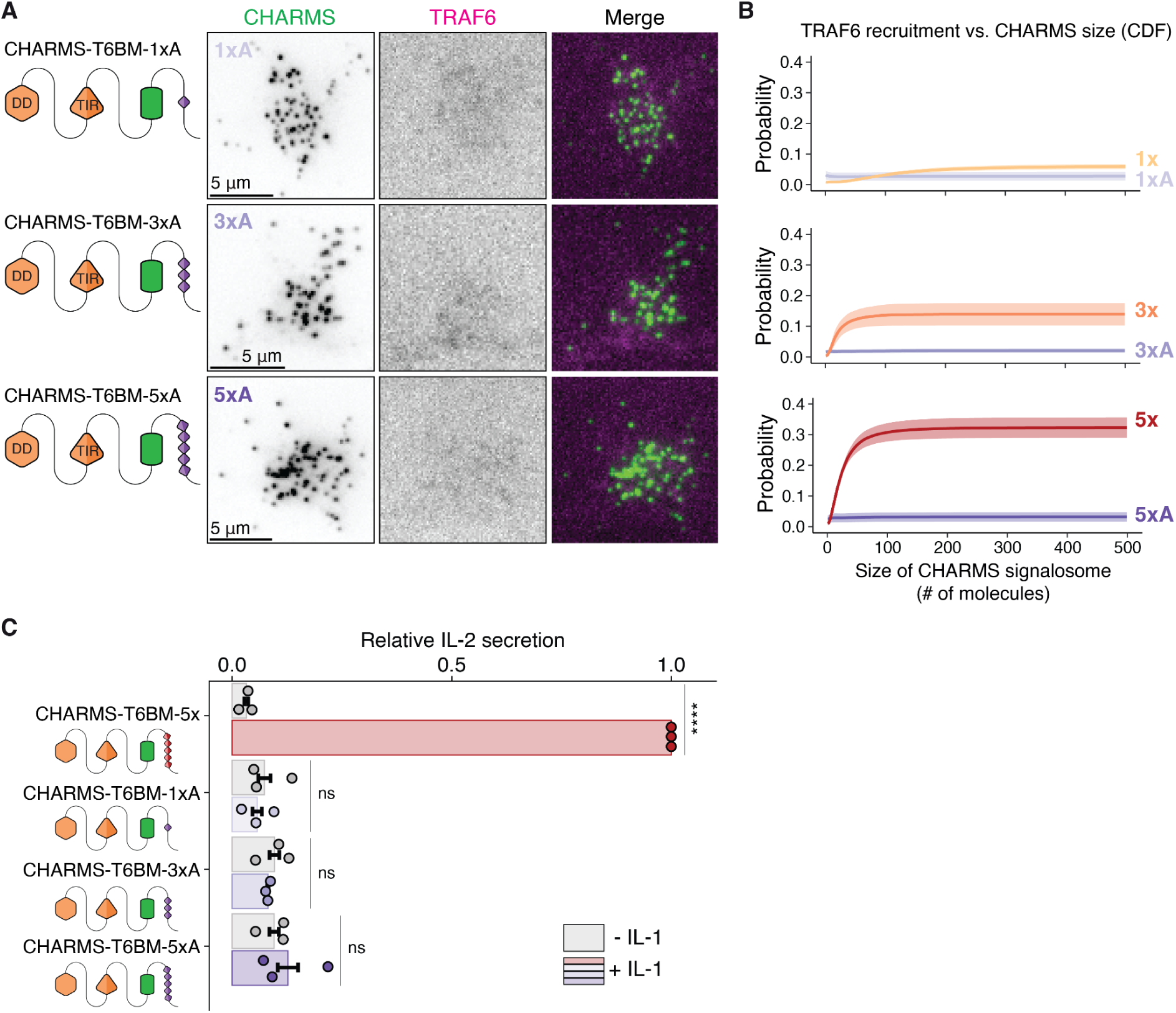
CHARMS constructs with a mutated TRAF6 binding motif are unable to signal. **(A)** We designed CHARMS with 1, 3 and 5 mutated T6BMs (PxAxxZ) - CHARMS-1xA (top), CHARMS-3xA (middle) and CHARMS-5xA (bottom), and reconstituted them into 3xKO cells. TIRF imaging confirmed all three CHARMS variants are defective in recruiting TRAF6. Scale bar, 5 µm. **(B)** The cumulative distribution function (CDF) of the probability of TRAF6 recruitment versus the size of CHARMS signalosomes with mutated T6BM compared with wild type T6BM. Shaded area indicates s.e.m. **(C)** IL-2 ELISA showing that the cells expressing CHARMS-1xA, 3xA and 5xA are unable to signal in response to IL-1. The IL-2 signal was normalized to the signal of the IL-1 stimulated EL4-MyD88-GFP/mScarlet-TRAF6 cells, referred to as WT cells in this experiment.

